# Structure, dynamics and evolution of the *Candida albicans* multi-drug resistance ABC transporter CDR1

**DOI:** 10.1101/2024.10.28.620768

**Authors:** Jeeeun Shin, Ruitao Jin, Barnabas Gall, Chacko Jobichen, Colin Jackson, Melanie Rug, Ben Corry, Joseph Brock

**Affiliations:** Australian National University, Research School of Biology, Division of Biochemistry and Biomedical Science; Australian National University, Research School of Chemistry; Australian National University, Centre for Advanced Microscopy

## Abstract

The pleiotropic drug resistance transporter Cdr1 from *Candida albicans* plays a crucial role in antifungal resistance. Here, we present high-resolution cryo-electron microscopy structures of *C. albicans* Cdr1 in multiple functional states: nucleotide-bound, substrate-bound, and apo forms. The 3.5 Å resolution structure of Cdr1 in complex with ATP and ADP reveals the molecular details of its asymmetric nucleotide-binding sites (NBS), with ATP bound to the deviant NBS1 and ADP to the canonical NBS2. Structures of Cdr1 bound to rhodamine 6G (3.4 Å) and Oregon Green 488 (3.5 Å) in complex with these nucleotides elucidate the pleiotropic substrate-binding pocket and highlight how nucleotide exchange drives conformational change required for transport. Additionally, we determined a 3.7 Å resolution structure of Cdr1 in the detergent LMNG without nucleotides, as well as a 3.5 Å resolution structure with nucleotides but no substrate, representing an apo state. We complemented these structural insights with molecular dynamics simulations to understand substrate binding dynamics, ancestral sequence reconstruction to trace the evolution of key functional motifs, and analysis of clinical isolates from sequence databases to identify potential resistance-associated variations. Comparison of these structures provides new insights into the conformational changes associated with the transport cycle of this asymmetric ABC transporter, revealing how ATP binding at the deviant NBS1 allosterically regulates the canonical NBS2, driving ATPase activity. This work significantly advances our understanding of the molecular mechanisms underlying multidrug resistance in pathogenic fungi and provides a structural and evolutionary framework for the rational design of Cdr1 inhibitors to combat antifungal resistance.

## Introduction

Untreated invasive candidiasis can lead to life-threatening complications despite the efficacy of topical treatments for milder cases. Immunocompromised individuals such as HIV/AIDS and COVID-19 patients in the healthcare facilities are considered the most vulnerable to invasive candidiasis and co-infection, with co-morbidity COVID-19 cases leading to mortality having recently been highlighted by the Centre for Disease Control and Prevention (CDC).^1^ Recent outbreaks of candidiasis in hospitals combined with an increasing number of clinical isolates exhibiting drug resistance around the globe signifies time for action. Recently, there has been an increasing report of multidrug resistance^2–6^ against antifungal azole type drugs such as fluconazole in the *Candida* species^3–9^, with *Candida auris* and *Candida albicans* being classified as critical pathogens in the fungal priority report of the World Health Organization (WHO).^7^ Although *C. albicans* is a common pathogen, primarily threatening immunocompromised individuals, *C. auris* has been designated an “urgent threat” by the CDC.^1^, Invasive *Candida spp.* infections currently impact 1.6 million people annually, leading to 995,000 deaths (63.6%). ^8^

Azoles are the most commonly used class of antifungals used to treat candidiasis. Their target is the cytochrome P450 lanosterol 14-α-demethylase (Erg11), that when inhibited blocks ergosterol biosynthesis and leads to the build up of a toxic sterol produced by the Δ-5,6-desaturase (Erg3).^9^ The multidrug resistance (MDR) against azole drugs in *Candida* species arise from a variety of mechanisms, including mutations affecting drug target Erg11, alterations in the ergosterol pathway, increase in sphingolipid biosynthesis, and overexpression of membrane transporters associated with the efflux of azole drugs. ^9–12^ While numerous membrane transporters are implicated in azole drug resistance, the upregulation of *CDR1* gene encoding *Candida* Drug Resistance Protein 1 (Cdr1) stands out as a predominant cause of efflux-mediated drug resistance in clinical *Candida* isolates.^4,9,11,13^ This observed upregulation frequently results from either an increase in gene copy number or the acquisition of gain-of-function mutations in the transcription factors responsible for Cdr1 overexpression. ^11^

Cdr1 protein exemplifies the structural and functional asymmetry of the pleiotropic drug resistance (PDR) subfamily of type V ATP-binding cassette (ABC) transporters^14^, known for their ability to efflux a diverse range of amphiphilic and hydrophobic substrates, including lipids, sterols, xenobiotics and rhodamine dye compounds.^15–18^ Much of the knowledge of this family comes from studies on its founding member, Pdr5 from *S cerevisiae,* that shares ∼55% sequence identity with Cdr1.^19^ Other fungal PDR family ABC transporters, including Pdr18, as well as the distantly related human ABCA members, have been implicated in sterol and lipid transport as primary function and all shown to be associated with drug resistance.^20–23^ The ability of Cdr1 to transport phospholipids has been postulated in the past and it has also been referred to as ‘floppase’^24^, however, the endogenous substrate of Cdr1 is still not known definitively. While the *CDR1* gene is not essential for cell viability, its deletion results in a drug-sensitive phenotype.^13^

The fungal PDR subfamily evolved through an ancestral gene duplication event, resulting in marked structural and functional asymmetry between the two halves of its members, in contrast to the dimeric architecture of homologous mammalian transporters^25^. Thus, the gene contains a nucleotide binding domain (NBD) and transmembrane domain (TMD), connected end to end with another C-terminal NBD-TMD. Recent reviews of the wealth of literature that has studied the structure and mechanistic principles of Cdr1^15^ in addition to *S. cerevisiae* Pdr5^19^, provide an excellent overview of the implications this has for the mechanism of transport. In summary, the two TMDs form a “Λ” shape in the inward-facing (IF) confirmation within the membrane that forms a cleft where substrates bind, with the NBDs located in the cytoplasm. Transport occurs when nucleotides bind and are subsequently hydrolysed at one of the NBDs. This provides energy that forces the NBDs to dimerise, squeezing the substrate binding pocket in a peristaltic pump motion that extrudes the substrate into the outer leaflet of the membrane.

The asymmetric structure of the transporter means that the two halves have different functions, with each nucleotide-binding site being quite distinct. While both NBDs are still capable of binding nucleotides, NBS1 (N-terminal) exhibits degenerate or ”deviant” motifs and lacks ATPase activity, while NBS2 maintains canonical sequences required for ATP hydrolysis. The canonical motifs in NBS2 include: the Walker A motif (GXXGXGKS/T) for ATP binding; Walker B motif (hhhhDE, where h represents a hydrophobic residue) essential for ATP hydrolysis; Q-loop involved in Mg2+ coordination and NBD-to-TMD communication; D-loop mediating NBD dimerization and inter-NBS communication; and H-loop containing the catalytic histidine. In contrast, the deviant NBS1 contains several key substitutions: a modified Walker A (GNVGKS/T), Walker B (hhhhDD), and signature sequence (L<u>NVE</u>Q). Notably, the D-loop sequence (GLD) remains conserved in both NBDs, highlighting its critical role in inter-NBS regulation. The signature motif or C-loop also interacts with ATP’s γ-phosphate in the opposite NBD upon dimerization of the two domains. Therefore, it is canonical in NBS1 (VSGGE) and mutated in NBS2 (LNVEQ)^15^ (**Table 1**).

**Table 1:** Comparison of NBD motif sequences. Underlined residues indicate unique substitution observed in the degenerative NBD motifs when compared to common consensus sequence.

| Motif | NBD1 (degenerative) |  | NBD2 (canonical) |  |
| --- | --- | --- | --- | --- |
|  | AA position | Seq | Seq | AA position |
| Walker A | 192 – 195 | GCST | GKTT | 900 - 903 |
| Walker B | 326 – 328 | <u>W</u>DN | LDE | 1025 - 1027 |
| Q-loop | 238 | E | Q | 942 |
| Signature/C-loop | 1001 -1005 | LNVEQ | VSGGE | 303-307 |
| D-loop | 1031-1033 | GLD | GLD | 332 - 334 |

ATP hydrolysis is crucial for Cdr1’s function in expelling a wide range of xenobiotics. However, it is noteworthy that the PDR subfamily exhibits “uncoupled” transport, characterised by high basal ATPase activity that is not stimulated by the presence of substrate, distinguishing them from most other ABC transporter families. The recent cryo-EM structures of Pdr5^26^ revealed that ATP binding at NBS1 is maintained in both inward-facing and orthovanadate-trapped outward-facing conformations, while NBS2 undergoes cycles of ATP binding and hydrolysis. This has provided a structural basis for this behaviour and enabled the proposal of a transport mechanism. ^26,27^ In this model, ATP is bound in the deviant nucleotide-binding site (NBS1) and ADP in the canonical site (NBS2), regardless of the presence of a substrate. The exchange of ADP for ATP within NBS2 is hypothesised to catalyse the dimerization of the opposing NBSs, triggering a conformational change to the outward-facing (OF) state that facilitates substrate release to the outer leaflet/extracellular space. ATP hydrolysis at the canonical site then provides the energy required to return the transporter to the inward-facing state (**Figure 1**). Thus, the presence of substrate does not stimulate ATPase activity, but it is required for transport to occur. A crucial observation in understanding how this allosteric regulation is mediated was the identification of a conserved linker domain that connect NBD1-TMD1 and contains a nucleotide sensing motif within the structure^37^ (^800^KQKG**E**IL^806^ or ^790^<u>M</u>QKG**E**I<u>V</u>^796^ in Pdr5 and Cdr1, respectively), with an acidic residue (in this case glutamate, bold) being essential. This motif contacts the bound ATP in both outward and inward facing conformations but is disordered in apo structures. While mutations within this domain decrease V_max,_ they have little effect on K_m_ of ATPase activity at NBS2, suggesting a role in allosteric regulation.^26,27^ ATP binding at NBS1 may also induce conformational changes that are transmitted to NBS2 via the C- and D-loops during dimerisation, modulating its ATPase activity.^15,26,27^ Phylogenetic studies of fungal ABC transporters indicate that the PDR family is the least conserved due to rapid evolution following the divergence of major fungal lineages, driven by multiple gene duplications and losses, which complicates the identification of orthologous PDR proteins across different species.^25,28^ To date, there has been no extensive ancestral reconstruction of the PDR family to reveal Cdr1’s evolutionary trajectory, nor has a large-scale bioinformatic analysis of global Cdr1 sequence variation been undertaken.

**Figure 1.**
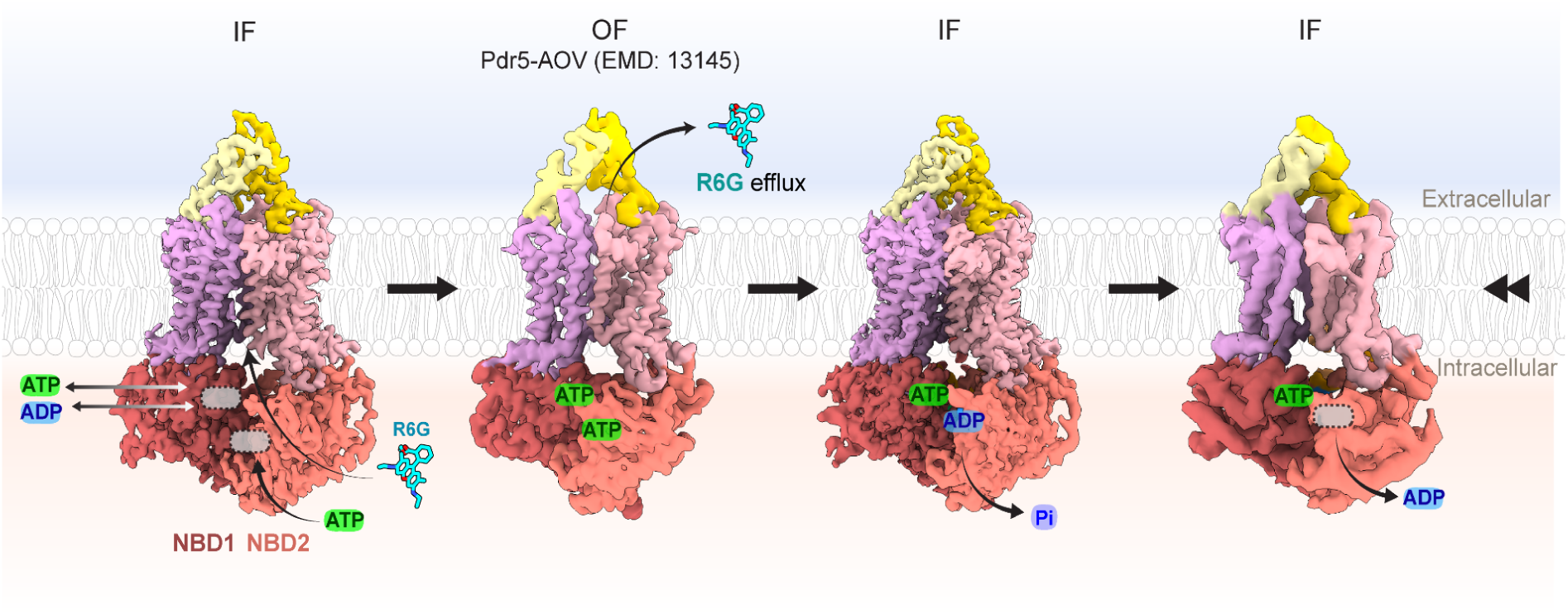
The full mechanistic cycle of Cdr1, modelled on current data of Cdr1 and outward-open conformation of its homolog Pdr5 (EMD:13145).^26^. The transport cycle begins with detergent, lipid, or an endogenous substrate entering the transmembrane domain (TMD) via an intracellular lateral entrance while the transporter rests in an inward-facing state (IF). Our results are consistent with the hypothesis that the deviant NBD1 acts as a sensor for ATP concentration in the cell^39^ and may bind ADP under nutrient limited settings. The binding and release of ATP or ADP in NBD1 are represented by a two-way arrow. ATP binding to the degenerative NBD1 brings the NBDs closer together in this conformation. Followed byATP binding at the canonical NBD2, the NBDs dimerise and transition to an outward-facing (OF) conformation, expelling the substrate (represented as R6G in this case). Hydrolysis of ATP at NBD2 then provides energy to return to the inward-facing state (IF). Pi is released, leading to the dissociation of ADP from NBD2, while ATP remains bound to NBD1.During this stage, substrate can access the TMD in an uncoupled fashion. The mechanical cycle returns to the primary substrate and nucleotide-accepting stage, as shown by the rewind symbol. For the diagram,the density-modified Cdr1-R6G represents the first inward-facing (IF) map, while the nucleotide-bound Cdr1-ATP is used to demonstrate post ATP hydrolysis IF. To illustrate the full mechanistic cycle, the outward-facing (OF) cryo-EM map of Cdr1’s homolog Pdr5 (EMD:13145) was included. Trajectory 1 from the 3DV map shows ATP binding in NBD1. The cryo-EM maps in the schematic were contoured at σ = 0.9–6.

## Results and discussion

In this work, we elucidated the cryo-EM structures of detergent-reconstituted Cdr1 with Oregon Green 488 (Cdr1-OG488) and Rhodamine 6G (Cd1-R6G) in the presence of ATP and ADP, bound at NBS1 and 2, respectively. We also observed a nucleotide-bound state without substrate (Cdr1-ATP/ADP) and unexpectedly identified the Lauryl Maltose Neopentyl Glycol (Cdr1-LMNG) detergent-bound state in the absence of nucleotides, despite including 100 μM of the azole drug Fluconazole during grid preparation. This may provide support for the proposed role of Cdr1 as an endogenous lipid transporter. ^24^ Recently, Peng and colleagues^29^ published several other structures of CDR1, providing detailed insight into the substrate recognition of fluconazole and the inhibitory mechanism behind milbemycin oxime. These structures, like the complex with LMNG presented in this study, are in the absence of nucleotides bound at the NBSs. Our study is therefore complementary, since we also were able to resolve the ternary complexes of OG488 and R6G together with ATP and ADP, bound at the deviant and canonical NBS, respectively. The high-resolution visualisation of this state, hypothesised to be the penultimate step preceding transition to the outward confirmation, is essential to understanding the unique allosteric modulation of the transporter. To this end, we have studied the binding modes of substrates and the comparative affinity of ADP/ATP at the deviant NBS1 within the ternary complexes using atomistic molecular dynamics (MD) simulations.

To further understand the structure-function relationship of Cdr1, we have also performed a thorough bioinformatic analysis of extant and ancestral sequence variation. Although significant sequence homology between Cdr1 and Pdr5 suggests a close evolutionary link with MDR traits, recent studies combining Cdr1 inhibitors and antifungals have selected for resistance mutations within regions of evolutionary divergence, highlighting their role in drug resistance mechanisms. ^30,31^ Although gain of function mutations within the Tac1/Tac1B transcription factor locus is an established mechanism for increasing Cdr1 expression in MDR *Candida spp.* clinical isolates^11,32^, sequence variations within Cdr1 associated with MDR in these samples are not studied-extensively. We hypothesised that the selection pressure from antifungal medications may have also induced sequence variation to evolve in the global population that would support the MDR phenotype. While large datasets of *Candida spp.* diversity have been publicly deposited^3,33,34^, these have not been extensively studied for clinically significant mutations in Cdr1. We sought to investigate the global sequence variation in *C. albicans* Cdr1 by downloading all *C. albicans* accession contigs that mapped to our reference sequence from the sequence read archive (SRA) using the LOGAN database.^35^ We also sought to understand the evolution of the gene using ancestral sequence reconstructions to identify how conserved and mechanistically important regions of the transporter have changed over evolutionary time.

### Asymmetric architecture of CDR1 as a full-length ABC transporter

We determined the cryo-EM structures of *C.albicans* Cdr1 protein using purified sample solubilized with LMNG detergent (**Figure 2A)**. We report the structures of Cdr1 in four different functional states, viz; no ATP (Cdr1-LMNG), only ATP/ADP nucleotides (Cdr1-ATP/ADP), as well as Rhodamine 6g (Cdr1-R6g) and Oregon green 488 (Cdr1-OG488) with ATP/ADP bound within NBD1 and NBD2, respectively. The datasets were collected using a JEOL CryoARM 200 fitted with a DE64 camera. The final cryo-EM map resolutions ranged between 3.45 – 3.73 Å and this resolution range was sufficient to resolve side chains, especially for the helices found in TMD where binding of ligands was observed (**Figure 2B**). Our cryo-EM structures of Cdr1 agree with well-established characteristics of an asymmetric full-transporter architecture in type V fold. ^14^

**Figure 2.**
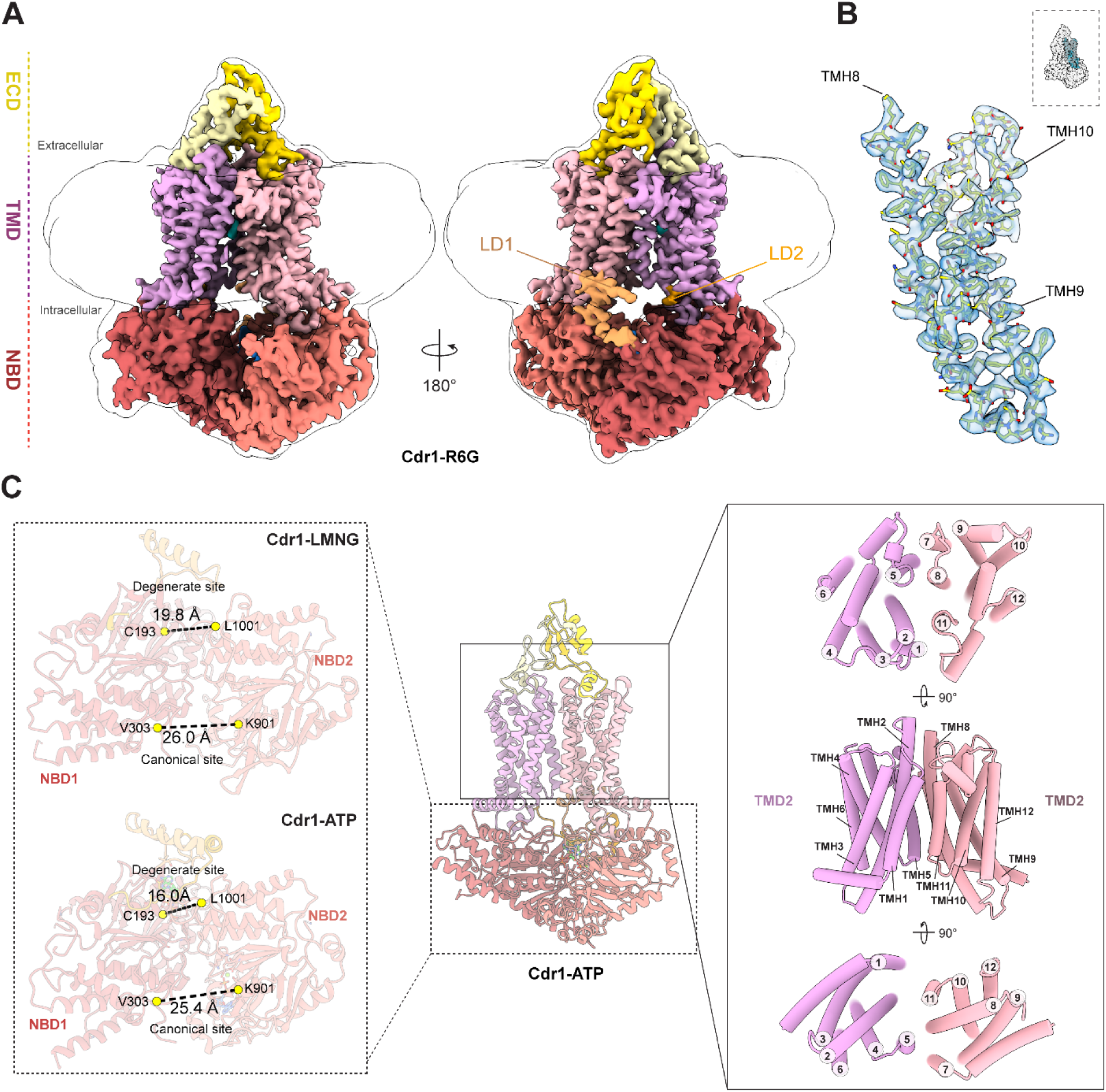
Cryo-EM structure of Cdr1 in the inward-open state. **(A)** Cryo-EM map of Rhodamine 6G bound Cdr1, shown in the inward-open conformation (σ = 4.3) . A 3σ Gaussian-filtered map (white) overlays the Cdr1-R6G density, with domain organization coloured accordingly. **(B)** Modelled TMH 8–10 (residues 1222–1337) of Cdr1-R6G fit into the Cryo-EM density (σ = 1.4). **(C)** Distances between Cα positions of the signature motifs and Walker A residues were measured, specifically from C193 in the Walker A motif of NBD1 to V304 near the signature motif of NBD2, as well as between K901 in NBD1 and L1001 in NBD2. The middle panel shows the side view of refined Cdr1-ATP structure in the inward-open state. The right panel provides top, side, and bottom views of the transmembrane α-helices (TH1–12) as cylinders.

The full transporter could be separated into two main domains, N- and C-terminal domains, where each domain consists of three distinctive sub-domains: nucleotide binding domain (NBD), transmembrane domain (TMD) and extracellular domain (ECD). Like the recent Pdr5 structure, Cdr1 shows both TMDs and NBDs forming a pseudo dimer. ^26^ The inter-residue distance between C193 of the Walker A motif in NBD1 and L1003 of the signature motif in NBD2, and the distance between K901 of the Walker A motif in NBD2 and V304 of the signature motif in NBD1 were used to quantify relative motion of NBD2 towards NBD1. Binding of nucleotides led to an average decrease of 3.56 ± 0.30 Å in the Cα-Cα distance between the Walker A motif in NBD1 and the signature motif in NBD2 (Cdr1-ATP/ADP binary complex and Cdr1-OG488/R6G ternary complexes) relative to the nucleotide-free condition (Cdr1-LMNG) (**Figure. 2C**) (Supplementary Table 2). Conversely, the distance between Walker A in NBD2 and the NBD1 signature motif showed a smaller change upon nucleotide binding. Where the distance was 0.78 ± 0.31 Å lower than in absence of nucleotide (Supplementary Table 2). Each TMD, namely the N-terminal TMD1 and the C-terminal TMD2, comprises six transmembrane α-helices (TMHs). While the corresponding TMHs from TMD1 and TMD2 are structurally similar, they differ slightly in their positional arrangement, contributing to the overall asymmetry of the structure (**Figure. 2C)**. Particularly notable, with the close arrangement of TMH1-3 in TMD1 being less prominent in TMH6-9 of TMD2 when viewed from the outer leaflet. Additionally, from the intracellular region, TMH2-3 and TMH6 of TMD1 show distinct positional differences compared to their counterparts TMH8-9 and TMH12, in TMD2 (**Figure 2C**).

### Degenerative and canonical nucleotide binding domains of Cdr1

The canonical Walker A motif contains a conserved lysine residue whose ε-amino group coordinates with ATP’s β- and γ-phosphates, stabilizing nucleotide binding. In the deviant NBD1, this lysine is replaced by cysteine, which cannot provide equivalent ATP stabilisation. The most significant catalytic impairment in NBD1 stems from the Walker B glutamate to asparagine substitution, as this glutamate is crucial for polarizing water for nucleophilic attack on ATP’s γ-phosphate. The catalytic impairment is further compounded by the substitution of the H-loop histidine with tyrosine in NBD1, which prevents proper nucleophile formation. Our cryo-EM structure reveals that this tyrosine’s hydroxyl group forms a hydrogen bond with the γ-phosphate, suggesting it functions to stabilize rather than hydrolyse ATP. Comparing the NBDs of Cdr1 and Pdr5 reveals subtle differences in nucleotide interactions. While T914 in Cdr1’s NBD2 forms a non-essential hydrogen bond with adenine, the equivalent M924 in Pdr5 shows weaker interaction. The linker domain (LD) structure is conserved between both proteins, with residues E794 and I795 stabilizing ATP binding within NBD1 (**Figure 3A**). Despite the resolution making unequivocal identification difficult, ATP was modelled in NBD1 and ADP with Mg^2+^ in the NBD2 site (**Figure 3A-B**). This is supported by previous observations of Pdr5.^26^

**Figure 3.**
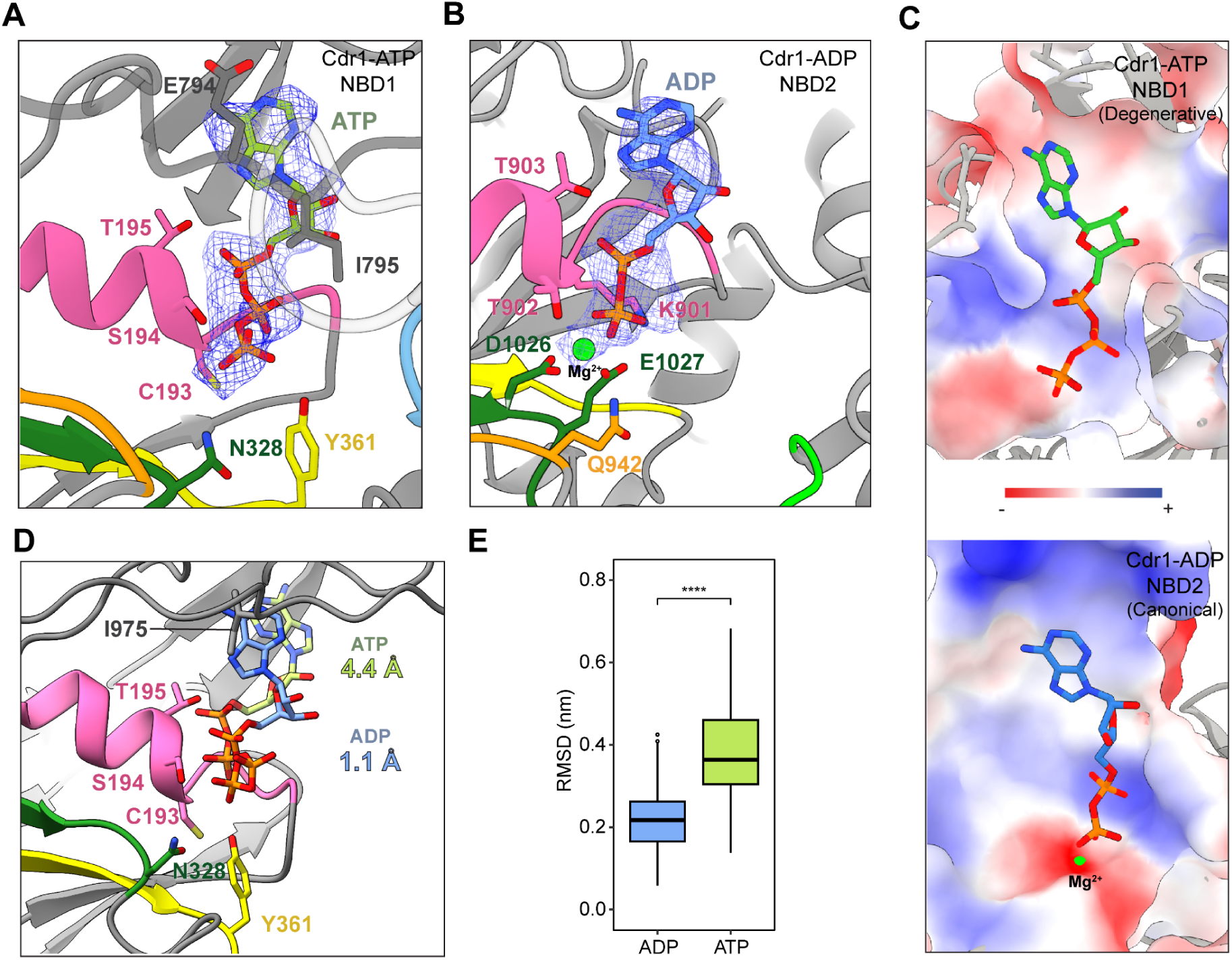
Nucleotide binding in Cdr1’s NBDs. (A) Cartoon representation of Cdr1-ATP structure with NBD1 in the ATP-bound state. (B) ADP-bound state, including Mg²⁺ ion (green sphere). Cryo-EM density for ATP and ADP+Mg2+ (σ = 3.5) are represented as blue mesh. Mg²⁺ ion (green sphere), Walker motifs A (pink) and B (forest green), along with the Q-loop (orange), H-loop (yellow), D-loop (lime green), and signature motif (sky blue), are highlighted. The ATP-sensing LD2 loop interacting with ATP in NBD1 is depicted in dark grey. All residues interacting with nucleotide or the Mg²⁺ ion is shown in stick representation. (C) ATP bound within the degenerative NBD1 and ADP bound within the canonical NBD2 shown as electrostatic surfaces, indicating a qualitatively more negatively charged binding site in NBD1. This may favour dissociation of negatively charged nucleotides under certain conditions via charge repulsion. (D) Snapshot of NBD1 binding with ADP and ATP bound after 800ns MD simulation with individual RMSD value indicating the shift from cryo-EM model respectively. (E) Root mean square deviation (RMSD) profile of ADP (blue) and ATP (green) in MD simulation shows that the ADP molecule binds more stably at the NBD1 site.

Functional studies revealed that peptidisc-reconstituted Pdr5 shows significantly reduced ATPase activity compared to membrane-embedded protein, likely due to increased rigidity. ^26^ Unlike Pdr5, we could not capture the OF conformation of Cdr1 via the addition of orthovanadate or AMP-PMP using the same methods, potentially due to high conformational heterogeneity associated with solubilisation in detergent. Additional cryo-EM map reconstructions of the Cdr1-OG488 and Cdr-R6G ternary complexes with added nucleotide also supported the modelling of ATP and ADP within NBD1 and NBD2, respectively. This indicates complete hydrolysis of the added ATP (2 mM) within the ∼5 minute incubation with OG488 and R6G.

### Relative nucleotide binding affinities within NBS1 and implications for mechanism

Analysis of our cryo-EM structures supports the hypothesis that Cdr1 effluxes substrates via uncoupled ATP hydrolysis. To determine the relative stability of ATP versus ADP in NBD1, we performed atomistic molecular dynamics simulations of our Cdr1-ATP model in a POPC bilayer. Our triplicate 800 ns MD simulations revealed that ATP is less stable than ADP in the NBD1 binding site, with ATP showing a tendency to move out of the pocket (**Fig 3D-E**). This suggests ADP is the more stable ligand at this site in this conformation. This is supported by the observation that the NBS1 of Pdr5 is much less negatively charged than at NBS2. ^27^ Our structures also confirm this to be the case for Cdr1 (**Fig 3C**). Within our maps however, clearly supported modelling ATP, which appears to be more stable given the 2 mM concentration of ATP added ∼5 min before grid freezing. Although we expect this had been largely consumed by ATPase activity within NBS2 during that period, the dissociation constraint of ATP at NBS1 may be slower than that of NBS1. Hence, there may exist an ATP/ADP binding equilibrium within NBS1 *in vivo*, that is dependent on intracellular concentrations (**Figure 1**).

While our cryo-EM studies, alongside others, demonstrate that Cdr1 and Pdr5 are stable in the apo-state^26,29^, questions therefore remain about ATP dissociation kinetics from NBD1 vs NBD2. 3D variability analysis (3DVA) of the Cdr1-ATP dataset showed that 20% of particle stacks Cdr1 are NBD2 ADP dissociated states (ATP bound in NBD1, no density in NBD2). Throughout the trajectories, ATP density persisted in NBD1. This loss of ADP density from NBD2, indicating ADP release, triggered a relaxation movement in NBD1 (Supplementary Figure 4), allowing us to capture this turnover cycle step. Despite applying a 4Å resolution filter for 3DVA however, precise model building proved challenging. Differences in co-factor densities were sufficient, however, to definitively observe ADP density loss within NBD2. This observation confirms the established role of the degenerate NBD1 in supporting NBD2’s ATPase activity. ^15,19^ Our results suggest that NBD1 may be bound by ADP under physiological conditions of limited nutrients, and only activates the transporter activity by binding to ATP once its concentrations increase above a critical level. This may be essential for limiting wasteful ATP consumption by this type of uncoupled transporter. A previous study characterising Cdr1 transport activity using a whole-cell transport assay indicated that drug-efflux is energy dependent. ^36^ Since the K_m_ of Cdr1 has been measured to be ∼1 mM^37^, while the *C. albicans* intracellular concentration of ATP has been measured at ∼ 1 μΜ^38^, the transporter may also exist in a nucleotide free state for significant periods *in vivo*.

### Presence of detergent bound in substrate binding cavity could be evidence of “floppase”

The Cdr1-LMNG structure, in the absence of ATP and MgCl2, revealed a pronounced inward-open conformation with increased NBD separation. Although the sample was prepared with 100 μM fluconazole, we observed LMNG detergent binding within the pocket instead, supporting the hypothesis that Cdr1 may function as a lipid transporter. ^24^ We postulate that LMNG enters the TMD binding cavity during reconstitution. For comparison, the concentration of LMNG was 100X less concentrated than that used to obtain the fluconazole binary complex recently reported by Peng and colleagues and our grid preparation used shorter incubation periods. ^29^ We hypothesise that the bulky glyco-diosgenin (GDN) detergent used for solubilisation likely couldn’t enter the TMD during the extended incubation used in those studies. In the presence of ATP, we hypothesise this would trigger IF to OF conformational switching, expelling the detergent and allowing R6G or OG488 to be observed in the substrate binding cavity. Although this could be an artifact of our experimental design, LMNG biophysically resembles a polar lipid, and could represent a physiological binding mode involved in the hypothesised floppase activity of Cdr1 supported by previous studies. ^17,24,27^

### Ligand binding pocket of Cdr1

The substrate promiscuity of Cdr1 and PDR superfamily is widely established by extensive biochemical and mutagenesis studies. The fluorescent compound Rhodamine 6G (R6G) is widely used for efflux assays and the recent structure of Pdr5 revealed the R6G binding pocket within the TMD. ^26^ We hypothesised that Oregon green 488 (OG488), a fluorescent compound we rationally selected due to its shared structural characteristics with R6G, would also be a substrate of Cdr1. We subsequently confirmed the binding of both R6G and OG488 with microscale thermophoresis (MST), with the binding of R6G weaker than OG488 (Supplementary Figure. 6, Supplementary Table. 4). The binding of OG488 was further confirmed with fluorescence polarisation (FP), using free OG488 as tracer (Supplementary Figure 7). From our ligand bound structures, the residues interacting with ligands were found in α-helices TMH1, TMH2, TMH5, TMH8 and TMH11 (**Figure 4A**). Our cryo-EM structures with R6G show that it binds deep in the CaCDR1 cavity. Interestingly, while the pkA of the dye (predicted as 10.3 by MolGpKa^39^) would suggest the presence of an iminium protonated N, in our structure this is buried within a number of hydrophobic residues. We thus sought to determine if the neutral un-protonated or charged protonated form of the dye is most likely to occupy this position. We found that the dye remains in the binding site in either situation (**Figure 4C**), and is slightly more stable in the protonated form. However, we found that water molecules enter the cavity to solvate the charged ethylazanium group to allow it to remain within the hydrophobic pocket (**Figure 4D**).

**Figure 4.**
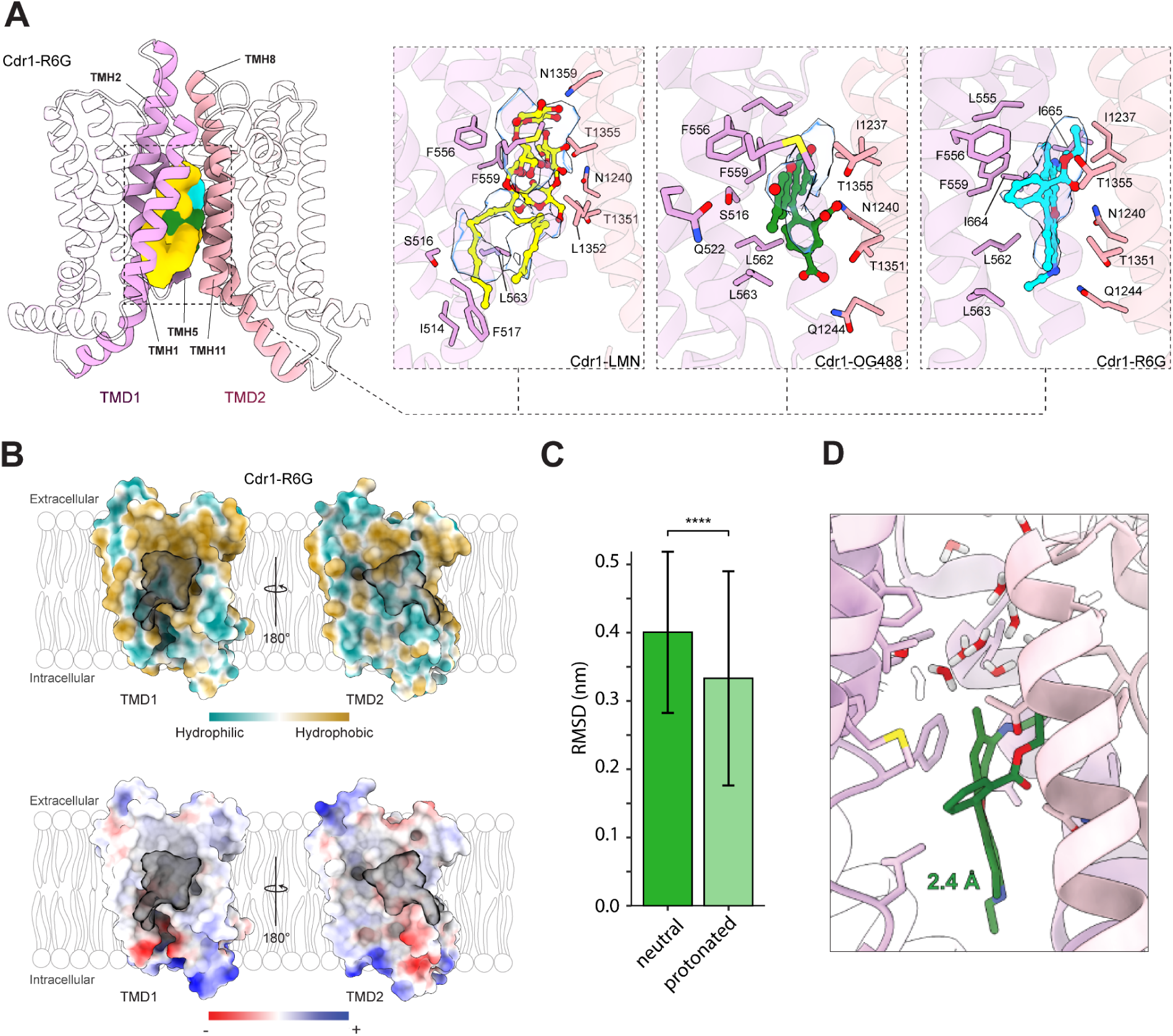
Ligand binding site of Cdr1. **(A)** Cartoon representation of the transmembrane domain (TMD) illustrating ligand-protein interactions in side view, highlighting TMH1, TMH2, TMH5, TMH8, and TMH11. Molecular surface representations of aligned LMNG (yellow), OG488 (forest green), and R6G (cyan) are shown at 4 Å resolution within the TMD cavity of the Cdr1-R6G model. Ligand-protein interactions are detailed, with interacting residues shown as ball-and-stick models. Cryo-EM density for each ligand is depicted as a transparent blue surface for LMNG (σ = 0.9), OG488 (σ = 0.8), and R6G (σ = 2.3). **(B)** Surface representation of TMD1 and TMD2 from lateral side view using Cdr1-R6G model. The surfaces are coloured to represent the hydrophobicity (top) and electrostatic potential (bottom). A combined molecular surface representation of ligands at 4 Å resolution occupying the TMD binding cavity is shown in transparent dark grey**. (C)** The averaged Root mean square deviation (RMSD) profile of neutral R6G (darker green) and protonated R6G (lighter green) in MD simulation. **(D)** Snapshot of protonated R6G at the end of 800 ns MD simulation in one replica, water molecules on top of R6G are shown.

### Analysis of fluconazole binding poses

Since we were unsuccessful in observing Fluconazole in the substrate binding site of Cdr1, we sought to identify the binding positions of Fluconazole using atomistic molecular dynamics simulations. We first docked the compound and chose the 5 best poses as starting coordinates for triplicate 800 ns MD simulations for each (a total of 15 simulations totalling 12 μs). We find the compound remains deep within the cavity in all cases but can adopt a range of different orientations/poses as indicated but the density surface (**Figure 5A**). These allow interactions with a wide range of cavity lining residues with the closest interactions identified in Fig 5B which are also conserved in the interactions formed with R6G compound. Most of these interactions are hydrophobic in nature, although hydrogen bonds can form to some amino acid protein residues. To better understand the range of positions we plot the likelihood of finding the compound at different z positions and at different RMSD from the modelled ligand position in the recent cryo-EM structure^29^ (**Figure 5C**). This shows that while some poses are very close to the model, there are a range of orientations found with different frequencies. To capture this diversity, we cluster the ligand positions (using a 3 Å cutoff) (**Figure 5D**) and find a number of poses shown in Fig 5E that represent the hot spots in Figure 5E (as well as multiple less populated clusters one of which overlaps the cryo-EM modelled position). These share a common location to the modelled pose in the cryo-EM structure but with different orientations of the fluconazole ‘arms’. Together this shows that while fluconazole binds stably within the cavity, it can adopt a range of orientations due to it forming multiple non-specific interactions with surrounding residues rather than specific hydrogen bonds. Although Peng and colleagues^29^ recently observed density modelled as a single conformation of Fluconazole using a very high (10 mM) concentration of the compound during grid preparation, we hypothesise multiple conformations are likely for this relatively small substrate under physiological concentrations.

**Figure 5.**
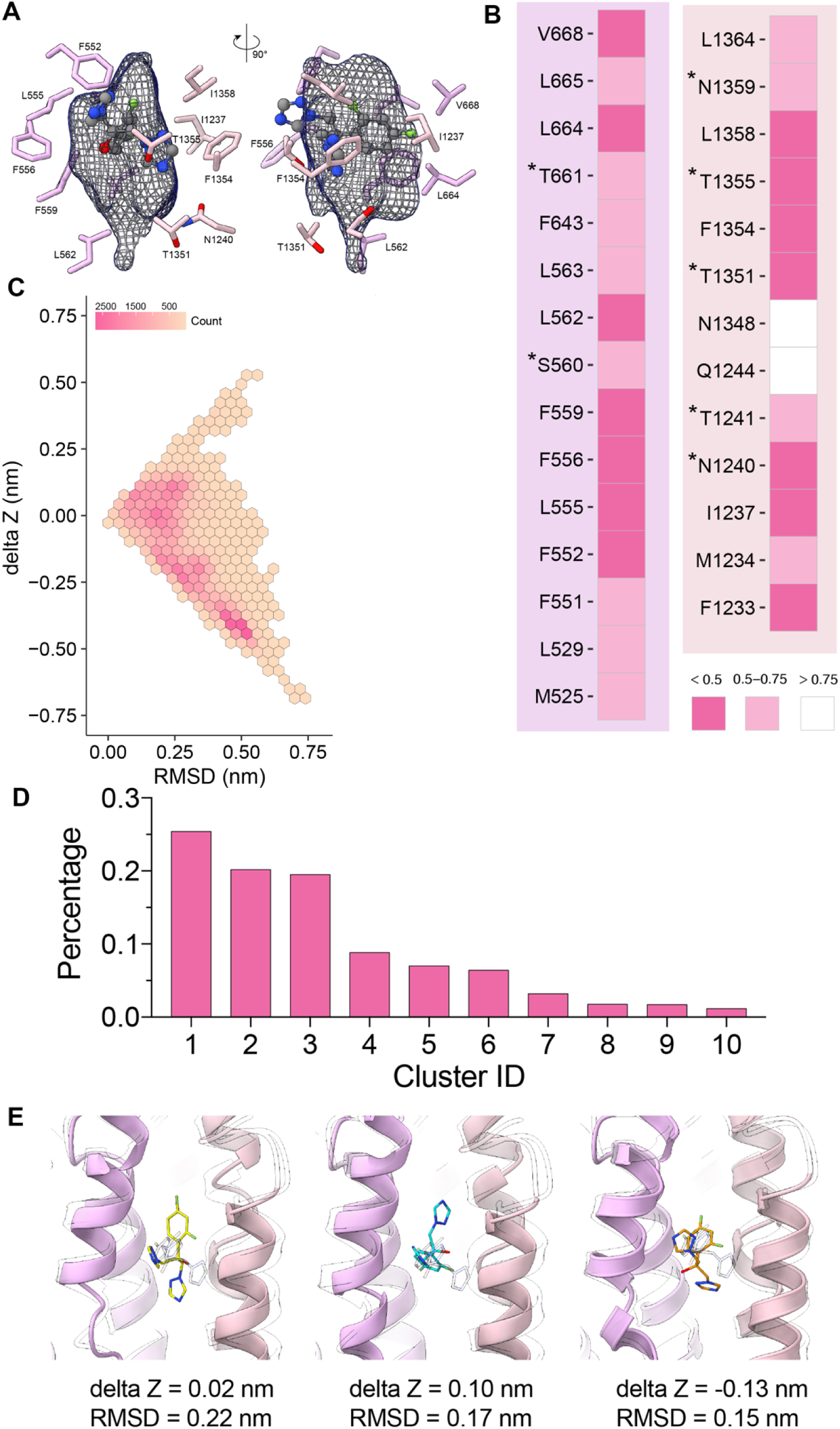
Investigation of Fluconazole binding modes using molecular dynamics simulations. **(A)** Averaged volumetric density of ligand Fluconazole using combined 12-µs-long MD trajectory in blue mesh with a resolution of 0.5 Å, the surrounding residues are potential contacting residues inside of transporter cavity. **(B)** Time-averaged minimal distances between residues lining the transporter cavity that could form interaction with Fluconazole. Residues with ∗ symbol can form hydrogen bonds with fluconazole according to MD trajectory. The colour gradient from darker to lighter indicates the distance ranging from smaller to bigger values. **(C)** 2D position distribution of Fluconazole plotted according to the RMSD of the centre-of-mass (COM) of fluconazole to the modelled fluconazole in previous cryo-EM structure, and the relative z-position of the centre-of-mass (COM) of Fluconazole, respectively. **(D)** The relative population of top 10 clusters identified from MD simulation in percentage using a cut-off of 3Å. (**E)** Three representative binding poses of fluconazole (whose carbon atoms are in yellow, blue and orange, respectively) identified from simulation in overlay with the modelled fluconazole in previous cryo-EM structure (carbon atoms in transparent grey).

### Insight into the changes within a protein sequence across evolution - Ancestral reconstruction

To understand the evolutionary history leading to CDR1, five replicates of maximum likelihood tree modelling were performed on a multiple sequence alignment (MSA) with members belonging to the Dikarya subkingdom. The MSA for phylogenetic inference shows complete conservation of the Walker A,B Q-loop and D-loop of both the degenerative and canonical Nuclear Binding Domains. We then assessed the progression of ancestral sequences from the last common ancestor of Ascomycota to *C. albicans* CDR1. We observed little divergence in the transmembrane helices 2,5,8 and 11. However, interestingly a double glycine is present in the common ancestor in place of alanine at positions 1346-7, suggesting increasing flexibility in TMH11, which has important entropic ramifications for ligand binding in nearby Threonines 1351,1355, which were observed to interact with Rhodamine. Furthermore, we observe position 661 originates from an alanine, evolving to a serine and then to a threonine, showing a polarisation of this position. The ancestral sequences 1-9 have N1240 and T1241 whereas 11-13 have G1240 and N1241, suggesting the asparagine, which interacts with Fluconazole in *C. albicans* CDR1, is important and conserved, but has undergone a reciprocal substitution (**Figure 6**).

**Figure 6:**
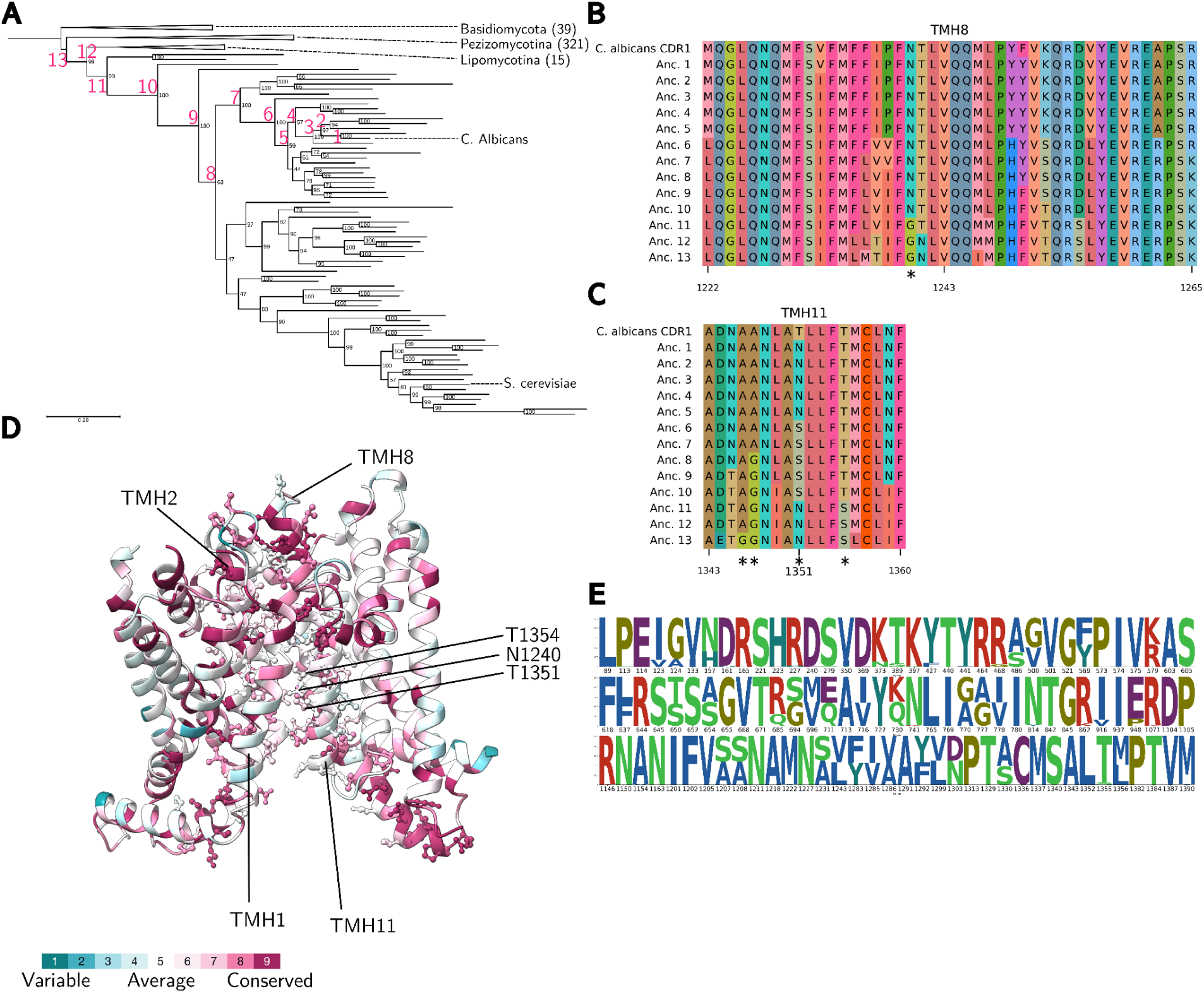
Phylogenetic analysis of CDR ABC transporters. **(A)** Maximum likelihood tree of representatives from the Dikarya CDR family. The tree is rooted between Basidiomycota and Ascomycota. Bootstrap support is indicated at each internal node. Pink numbers on internal tree nodes label ancestral sequences of *C. albicans* CDR1 shown in alignments B and C. **(B-C)** Alignments of transmembrane helices of C. albicans CDR1 and its ancestors: **(B)** helix 8 (positions 1222–1265) and **(C)** helix 11 (positions 1343–1360). **(D)** Consurf^40^ analysis of evolutionary conservation in the CDR1 transmembrane domain, calculated using the solved structure, curated multiple sequence alignment, and maximum likelihood tree. **(E)** Sequence logo^41^ of polymorphisms in CDR1 sourced from the LOGAN database. ^35^

### Critical Gating Residues in CaCdr1 Control Drug Efflux

The analysis of our structures of CaCdr1 revealed that three key residues - F673, L1364, and F1230 - are positioned at the interface between TMD1 and TMD2 near the outer leaflet of the membrane (**Figure. 7A**).^15^ This arrangement differs from the gating mechanism in Pdr5, where the equivalent residues (F683 and M1373) form a direct hydrophobic seal **(Figure. 7B,C)**. ^19^ While F673 and L1364 in CaCdr1 make contact with substrates, functional studies showed that their mutation to alanine did not significantly impact rhodamine 6G efflux. Instead, F1230 appears to play a critical role by projecting its phenyl ring into the substrate pathway, contacting both F673 and L1364 to modulate the exit channel. ^15^ The importance of these gating regions was further demonstrated through FK506 resistance studies. As reported by Tanabe and colleagues^31^, mutations near these gating residues, particularly F1235C (adjacent to F1230), were repeatedly isolated in FK506 resistance screens. The recurring nature of these mutations reinforced the functional significance of this region in controlling drug translocation. This distinct gating arrangement in Cdr1, while sharing architectural similarities with Pdr5, has evolved unique features with F1230 playing a more prominent role alongside F673 and L1364 in regulating substrate efflux. Understanding these critical gating elements provides important insights into the molecular mechanism of drug transport in this clinically relevant efflux pump.

**Figure 7.**
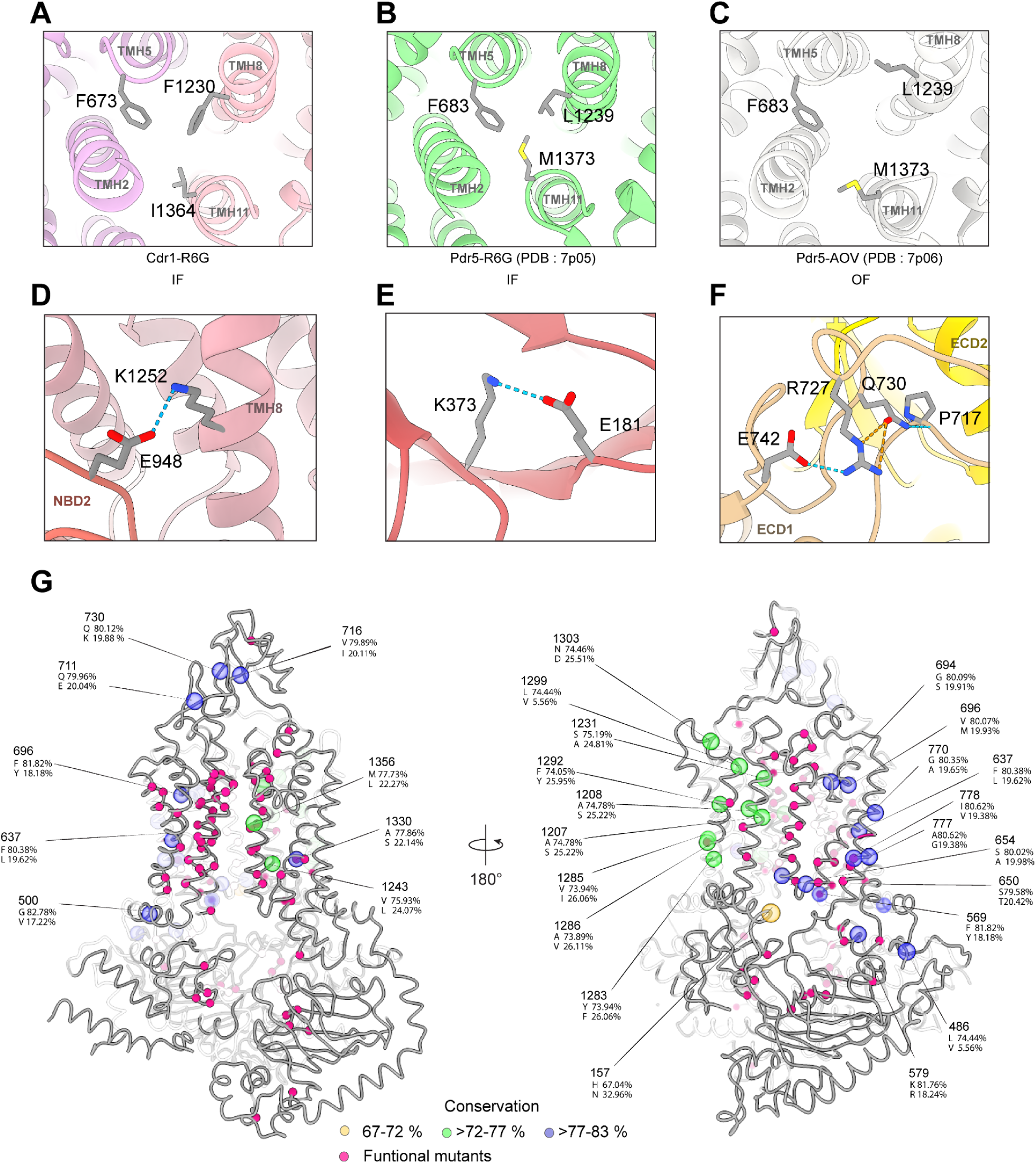
Gating residues and potential functional sites with high mutation rates are highlighted. Cartoon representations of Cdr1 and Pdr5 structures are shown: **(A)** Gating residues in the inward-facing (IF) conformation of R6G-bound Cdr1. **(B**) Gating residues of R6G-bound Pdr5 (PDB: 7p05) in IF conformation. **(C)** Pdr5 (PDB: 7p06) in outward-facing conformation. **(D)** Cytosolic view of the salt-bridge between E948 (NBD2) and K1252 (TMH8). **(E)** Cytosolic view of the salt-bridged between K323 and E181. **(F)** Extracellular view of the hydrogen bond network among residues E742, R727, Q730, and P717 in ECD1. **(G)** Residues previously studied for their impact on drug susceptibility, along with those showing <83% conservation, are mapped onto the cartoon of the inward-facing Cdr1-R6G model. The top number shows the residue position, with amino acid conservation percentages and codes displayed below. Mutated residues affecting Cdr1 function are highlighted in pink, while residues with 67–72%, >72–77%, and >77–83% conservation are shown in yellow, green, and blue, respectively. The Cα positions of these residues are depicted as spheres.

### Residues of potential functional relevance

By combining our cryo-EM data with a bioinformatic analysis of Cdr1 global sequence diversity within the LOGAN database (**Figure 6E**), we identified several residues that had not been previously investigated. The negatively charged E948 in NBD2, located near TMH8, forms a salt bridge with K1252, likely serving a structural role by linking the intra-domain junction between NBD2 and TMD2 (Figure. 7D). E948 is highly conserved at 98.54%, with only 1.46% represented by the variant P948. The E948P mutation compromises the salt bridge and losing this intra-domain interaction may adversely affect Cdr1 function. We also observed that K373 in NBD1 forms a salt bridge with E181 on an adjacent β-sheet extending from the Walker A motif (**Figure 7E**). This interaction appears to serve a structural role. Our CDR1 sequence variant analysis shows K373 is highly conserved (97.84%), with only 2.16% represented by the variant N373. Substitution of positively charged lysine with polar asparagine would likely form a weaker hydrogen bond, potentially affecting the Walker A motif’s positioning in NBD1. We identified another variant, Q730K, located in the ECD1 of Cdr1 (**Figure 7F**). Q730 shows 80.12% conservation, with the remaining 19.88% corresponding to K730. The hydrogen bond network involving P717, Q730, R727, and E742, which provides rigid structural integrity in Cdr1’s ECD1, is absent in its homolog Pdr5 in both IF and OF conformations. This difference suggests that this hydrogen bond network may contribute to Cdr1’s clinical MDR phenotype. The Q730K variant would disrupt this network, which may affect the efflux activity of Cdr1. Mapping residues with conservation scores below 83%, along with those previously investigated, showed that the majority of previous studies targeted ligand and nucleotide binding sites, whereas most variants within the LOGAN database were in ECD1 and non-binding site TMHs (**Figure 7G**). However, we did find several sites present in both datasets, such as F569, A1286, and A1330, although the substitutions we identified were not detected in prior mutatagenesis studies. ^15,31,42^ Notably, the N-terminal domain side contained more sequence variability, while the C-terminal side showed fewer variants and lower conservation (72–77%). To better understand the function and impact of these identified variants, further exploration of these residues and their variants could offer valuable insights into Cdr1’s role in multidrug resistance.

## Concluding remarks

In this study, we elucidated the structural and functional dynamics of the multidrug resistance transporter Cdr1 from *Candida albicans*, offering critical insights into its unique asymmetric architecture and mechanism of substrate transport. High-resolution cryo-EM structures of Cdr1 in various functional states, including nucleotide-bound and apo forms, revealed how ATP binding at the degenerate NBS1 site modulates ATP hydrolysis at the canonical NBS2, driving essential conformational changes required for substrate efflux. These structural insights underscore Cdr1’s ability to transport a broad range of substrates and highlight the roles of specific residues within its binding pocket and gating regions in mediating drug interactions and resistance.

Our findings demonstrate that the asymmetric nucleotide-binding sites function cooperatively, with ATP binding at the deviant NBS1 serving as a sensor for cellular energy levels that modulates ATPase activity at NBS2. Molecular dynamics simulations further support this by showing differential nucleotide stability at NBS1, suggesting a mechanism responsive to fluctuations in cellular ATP concentrations. This functional asymmetry also underpins Cdr1’s substrate promiscuity, as shown by LMNG binding, which suggests a putative endogenous role in lipid transport alongside its established xenobiotic efflux function.

Further mining of clinical sequence data using the LOGAN database revealed previously uncharacterized variants, such as Q730R and E181P. While these variations’ roles in multidrug resistance phenotypes remain to be elucidated, their presence in clinical isolates points to adaptive pathways that may work alongside transcriptional upregulation to enhance resistance. Ancestral sequence reconstruction revealed strong conservation of the Walker A, B, Q-loop, and D-loop motifs in both nucleotide-binding domains (NBDs), alongside key changes in transmembrane helices that may facilitate substrate binding flexibility. These include a double glycine-to-alanine shift in TMH11 and reciprocal asparagine substitutions near the drug-binding pocket, suggesting evolutionary tuning of substrate binding and efflux functions.

Together, our findings provide an advanced structural and evolutionary framework for understanding the molecular mechanisms of Cdr1 and open avenues for developing novel antifungal strategies to combat drug resistance in this clinically significant pathogen.

## Methods

### Expression and purification

Details of expression, detergent screening and purification of cdr1 were screened as described in protocol Drew and colleagues. ^43^ The *CDR1* gene from wild type, Fluconazole susceptible reference *Candida albicans* strain SC5314 was cloned into a pDDmTurq2, a derivative of pDDGFP-2 plasmid ^44^ which replaces yEGFP with mTurquoise2. Subsequently, mTuquoisw2 fusion Cdr1 was expressed and purified from S. cerevisiae FGY217 strain grown in -URA, 0.1 % glucose at 30 °C. Expression was induced at OD600 = 0.6 with addition of 2 % galactose in the medium and further incubated for 22 hours. Cell culture was centrifuged and resuspension in CRB buffer was lysed using heavy-duty cell disruptor at incremental pressures of 20, 22 and 25 kpsi and three passes of 30 kpsi at 4–15 °C. Cell lysate was collected by centrifuging at 10,000 x g at 4 °C for 10 min and collect the supernatant containing membranes. Membrane was collected by centrifuging at 150,000 x g at 4 °C for 1.5 hours. Subsequently, the pellet was resuspended in 6 mL of MRB per 1 L culture of membrane pellet and frozen for later purification. In this study, Fluorescence size exclusion chromatography (FSEC)^43^ was carried out to screen various detergents: Nonyl glucoside (NG), n-octyl-β-d-glucoside (OG), Octyl glucose neopentyl glycol (OGNG), n-Undecyl-β-D-Maltopyranoside (OGNG), n-dodecyl-β-D-maltoside (DDM) and Lauryl Maltose Neopentyl Glycol (LMNG). Membrane suspension was solubilised with 1% (w/v) of each detergent at volume of 150 μL, and centrifuged at 22,000 x g for 1 hr at 4 °C. The soluble crude membrane was isolated and filtered through a 0.22 μM filter. 100 μL of the filtered sample was injected onto the Superdex® 200 5/150 GL column equilibrated in DB buffer (20 mM Tris-HCl, 0.15 M NaCl, 0.03 % (w/v) DDM, pH 7.5). Each sample was eluted at a flow rate of 0.12 mL/min for 30 minutes and analysed for monodispersity using an inline fluorescence detector on a HPLC (Shimadzu).

For large scale purification of Cdr1, the *CDR1* gene from pDDmTurq2 plasmid was isolated and assembled into an episomal expression vector created using the MoClo yeast toolkit^45^ without an mTurquoise2 c-terminal tag (pGAL1-CDR1-twin-Strep/8xHis-tEN01). Subsequently, Cdr1 was expressed at a larger scale and the membrane was prepared as described above. Into membrane suspension, 1 % (v/v) LMNG was added and solubilized with gentle rotation for 1 hr and further centrifuged at 33,000 x g at 4 °C for 1 hr to remove aggregates. The detergent treated sample was filtered through 0.22 μM filter, then loaded onto 1mL Strep-tactin®XT 4Flow® FPLC column (IBA Lifesciences) using a peristaltic pump. Purification was performed as the manufacturer’s protocol. All fractions were pulled and dialysed in DB buffer (20 mM Tris-HCl, 0.15 M NaCl, 0.01 % (w/v) LMNG, 0.1 mM TCEP, pH 7.5) overnight and centrifuged at 10,000 x g at 4 °C for 1 hour, then filtered through a 0.22 μM filter. The filtered sample concentrated to a volume of 150 μL using a 100 kDa MW cut off Amicon® Ultra-15 Centrifugal Filter Units (Merck Millipore). Concentrated sample was further purified by size exclusion chromatography using a Superdex 200 10/300 GL (GE Healthcare, Chicago, IL, USA) column equilibrated in DB buffer without detergent (20 mM Tris-HCl, 0.15 M NaCl, 0.1 mM TCEP, pH 7.5).

### Microscale thermophoresis

Purified Cdr1 protein was labeled with Alexa Fluor 647 dye at 2-fold concentration of protein in HEPES (pH 7.56), 0.15 M NaCl, 0.1 mM TCEP, 0.001 % (w/v) LMNG. Free dye was removed by desalting using the PD-10 Desalting column (Cytiva). A labeled protein concentration range of 317 - 540 nM was used to perform MST using a red filter. A serial dilution of Rhodamine 6G and Oregon Green 488 was prepared and combined with the sample at a dilution factor of 1.7. The measurements were performed in standard capillaries (NanoTemper Technologies) using a LED source with 652 nm and 20% infrared-laser power at 25 °C. The results were fitted to Hill curve using OriginLabpro.

### Fluorescence polarization

Fluorescence polarization (FP) assay was performed by two-fold serial dilution of purified Cdr1 at concentration of 31.5 µM in DB buffer (20 mM Tris-HCl, 0.15 M NaCl, 0.01 % (w/v) LMNG, 0.1 mM TCEP, pH 7.5) with 1µM of OG488. FP measurement was acquired using EnVision® microplate reader (PerkinElmer) equipped with Optimized FITC FP Dual Emission Label (Excitation λ: 480 nm, Emission λ: 535 nm, Bandwidth = 40 nm). For the tracer, free Oregon green™ 488 (OG488 ; Excitation λ: 490 nm, Emission λ: 514 nm; Cayman Chemical) solubilised in FP assay buffer was used. All sample preparation and data acquisition were carried out at room temperature. The experiment was performed in duplicate.

### Cryo-EM grid preparation and data collection

A CRYO ARM 200 (JEOL) electron microscope equipped with an in-column Omega energy filter and DE64 detector was used for all cryo-EM data acquisition. For the Cdr1-LMNG data set, a 3 μL aliquot of the sample at concentration of 2.5 mg/mL containing 100 uM of fluconazole in DB was applied onto a glow-discharged UltraAuFoil grid (R1.2/1.3). The grid was blotted for 2 seconds at 5°C and 100% humidity using a Vitrobot Mark IV (Thermo Fisher Scientific, USA). A total of 3650 movies were collected by SerialEM software^46^ under counting mode at a magnification of 100,000× (pixel size 1.25Å) with a defocus range of -0.5 to -2.7 μm. Each movie comprised 30 frames recorded at a dose rate of 4.4e^−^/Å^2^ s^−1^.

For samples with ATP, purified sample at a concentration of 3 mg/mL was prepared with 100 μM of either R6G or OG88 in DB buffer (20 mM Tris-HCl, 0.15 M NaCl, 0.0005 % (w/v) DDM, pH 7.5). Subsequently, 3 μL of protein/ligand sample mixture was applied onto glow-discharged UltraAuFoil grids (R1.2/1.3). The Cdr1-ATP sample was prepared with 2mM ATP and 2 mM MgCl2, then incubated for 2 minutes on ice. The Cdr1-OG488 and Cdr1-R6G samples followed the same preparation, with an additional ∼5-minute incubation after adding 100 µM OG488 or R6G. All grids were blotted for 3 seconds at 4.5 °C at 95 % humidity using Vitrobot Mark IV (Thermo Fisher Scientific, USA). A total of 1165, 933 and 3825 movies were collected with SerialEM software^46^ for grids with ATP only, R6G and OG488, respectively. Counting mode at a magnification of 120,000× (pixel size 0.996Å) with a defocus range of -0.5 to -2.0 μm was applied. Each movie consisted of 34 frames recorded with a dose rate of 4.0e^−^/Å^2^ s^−1^. Detailed data collection parameters are shown in Supplementary Table 1.

### Cryo-EM data processing

The acquired data was processed using cryoSPARC-4.1.0. ^47^ The movie frames were aligned using Patch Motion Correction and Patch CTF estimation. After multiple rounds of 2D classification and class selection, the best 2D classes were used for template picking. 2D classes from template picker were then used as template for Topaz picking, except for the Cdr1-LMNG data set, where only the blob picker was used. ^48^

Particles were further 2D classified and multiclass *ab initio* and heterogenous refinement was performed. To improve the map, the signal from the micelle belt was subtracted and final 3D refinement was done using the subtracted particles except for the refinement of the Cdr1-R6G data set. A more detailed workflow of Cryo-EM map refinement is summarized in Supplementary Figures 2 & 3. All Cryo-EM maps of Cdr1 were viewed with UCSF ChimeraX. ^49^ The resolution of the refined electron density map of ATP only was at 3.54 Å, R6G at 3.45 Å, OG 488 3.49 Å and 3.73 Å for Cdr1-LMNG as estimated using the gold standard Fourier shell correlation (FSC) with a 0.143 threshold from cryoSPARC-4.1.0. ^47^ A 3D variability analysis of the Cdr1-ATP data, using a 4 Å resolution filter, identified three components that showed similar trajectories of conformational flexibility.

### Model refinement

All maps were density modified with resolve_cryo_em^50^ from Phenix.^51^ Predicted model from AlphaFold^52^ was docked into maps as an initial model and further models were further refined with a combination of realspace refinement in Phenix^53^, Coot^54^ and ISOLDE.^55^ Ligand models were obtained from PubMed as sdf file, converted into pdb format and subjected to eLBOW^56^ in Phenix. For all models, regions with poor Cα backbone density were subsequently deleted. MolProbity v4.5.1^57^ was used to evaluate the quality of all structures. All data and model statistics are reported in Supplementary Table 1 .

### Molecular dynamic simulation

The cryo-EM structures of CaCDR1 in complex with different nucleotides at NBD1/2 sites, and/or R6G dye molecules in transporter cavity were used to create the starting structures of simulation systems. The fluconazole binding structures were obtained using molecular docking. YASARA (version 20.8.23) was used to conduct the rigid docking using the R6G binding structure as the template and VINA program^58^, AMBER03 force field was used for charge assignment and determining which ligand bonds can rotate. Top-five docking poses were selected for another round of VINALS local search program to sample local ligand conformations, and refined complexes were then used to conduct simulations. Whole simulation systems were prepared using the CHARMM-GUI Membrane Builder server^59^, with N- and C-terminal residues patched as acetylated N terminus and methylamidated C terminus, respectively. The transmembrane region of protein was assigned by using the PPM v.2.0 server.^60^ The simulation cells comprised approximately 240,000 atoms, of which approximately 54,000 were water molecules, in a box of dimensions 12 × 12 × 17 nm^3^. The principal axis of the transmembrane domain was aligned along the z axis of box. TIP3P water parameters were used to solvate all systems. About 350 POPC lipids were contained in the lipid bilayer, sufficient Na^+^ and Cl^−^ ions were introduced by replacement of water molecules to bring the systems to an electrically neutral state at an ionic strength of 0.15 M. All simulations were performed using the GPU-accelerated GROMACS software package (version 2023.1). ^61^ CGenff program was used to parameterize the small ligand^62^ and CHARMM36m force field with CMAP corrections^63^ was used to parameterize all interactions within the systems.

Steepest descent energy minimization with tolerance of 1,000 kJ mol^−1^ nm^−1^ was followed by six sequential steps of equilibration with a gradual decrease in the restraining force applied to different components. The LINCS algorithm^64^ was applied for resetting constraints on covalent bonds to hydrogen atoms, which allowed 2 fs time steps for MD integration during the entire simulation. The particle-mesh Ewald algorithm^65^ was used for calculating electrostatic interactions within a cut-off of 12 Å, with the Verlet grid cut-off-scheme^66^ applied for neighbour searching, using an update frequency of 20 and a cut-off distance of 12 Å for short-range neighbours. A 12 Å cut-off was applied to account for van der Waals interactions, using a smooth switching function starting at 1.0 nm. Periodic boundary conditions were utilised in all directions. During the equilibration stages, the temperature was maintained at 303.15 K using a Berendsen-thermostat^67^ with a time constant of 1.0 ps. Protein/ligands, membrane and ion-water groups were treated independently to increase accuracy. The pressure was maintained at 1.0 bar by isotropic application of a Berendsen-barostat^67^, with a time constant of 5.0 ps. During production molecular dynamics, the temperature was maintained at 303.15 K using a Hose-Hover-thermostat^68^ with a time constant of 1.0 ps, and the pressure maintained at 1.0 bar using the Parrinello–Rahman-barostat^69^ semi-isotropically, with a time constant of 5.0 ps and compressibility of 4.5 × 10^−5^ bar^−1^. All different systems were run in 3 replicates, respectively, for 800 ns in production MD, together a total of 21.6 µs long simulations were conducted [Table M1].

**Table M1.** Summary of Molecular Dynamics Simulations.

| Model | Structure | Ligand in Cavity | NBD1 | NBD2 | Simulation length |
| --- | --- | --- | --- | --- | --- |
| cryo-EM | Cdr1-ATP | - | ATP | ADP | 800 ns × 3 replicas |
| cryo-EM | Cdr1-ATP | - | ADP | ADP | 800 ns × 3 replicas |
| cryo-EM | Cdr1-R6G | neutral Rhodamine 6g | ATP | ADP | 800 ns × 3 replicas |
| cryo-EM | Cdr1-R6G | protonated Rhodamine 6g | ATP | ADP | 800 ns × 3 replicas |
| VINA docked | Cdr1-R6G | fluconazole pose 1 | ATP | ADP | 800 ns × 3 replicas |
| VINA docked | Cdr1-R6G | fluconazole pose 2 | ATP | ADP | 800 ns × 3 replicas |
| VINA docked | Cdr1-R6G | fluconazole pose 3 | ATP | ADP | 800 ns × 3 replicas |
| VINA docked | Cdr1-R6G | fluconazole pose 4 | ATP | ADP | 800 ns × 3 replicas |
| VINA docked | Cdr1-R6G | fluconazole pose 5 | ATP | ADP | 800 ns × 3 replicas |

### Phylogenetic reconstruction

A sequence dataset was curated by blasting the *C. albicans* CDR1 sequence (Uniprot accession: XP_723209) in NCBI and JGI MycoCosm, as well as incorporating the IPR010929 CDR ABC transporter InterPro database, resulting in 5,717 homologous sequences. All species within the dataset were of the Dikarya subkingdom. These were then filtered by length falling outside two standard deviations, and subsequently clustered by CD-HIT at 0.8 identity. A HMM was constructed within each cluster, and the most representative sequence (highest bitscore) was selected from the cluster, resulting in the MSA for subsequent phylogenetic tree modelling. Three initial clades were removed, which were likely paralogs upon comparison to the species tree. The final MSA contained 263 sequences, incorporating Basidiomycota and Ascomycota. The MSA was trimmed to remove disordered regions at the N and C terminus that did not align well. The LG model was used to construct the maximum likelihood tree and five replicates were performed (Table M2). Ancestral sequences were reconstructed at each node by the empirical Bayesian method implemented in iqtree2. Indel events were also modelled using a Jukes-Cantor model of evolution. The tree was rooted between Basidiomycota and Ascomycota.

**Table M2:** Statistics across tree replicates:

| Tree | logL | deltaL | bp-RELL | p-KH | p-SH | p-KH | p-SH |
| --- | --- | --- | --- | --- | --- | --- | --- |
| 1 | -<br>333920.30<br>33 | 0.27342 | 0.191 | 0.49 | 0.856 | 0.192 | 0.545 |
| 2 | -<br>333923.66<br>42 | 3.6343 | 0.157 | 0.465 | 0.842 | 0.158 | 0.484 |
| 3 | -<br>333934.18<br>6 | 14.156 | 0.168 | 0.4 | 0.714 | 0.168 | 0.428 |
| 4 | -<br>333920.02<br>98 | 0 | 0.257 | 0.51 | 0.257 | 0.257 | 0.548 |
| 5 | -<br>333948.01<br>82 | 27.988 | 0.227 | 0.368 | 0.532 | 0.226 | 0.361 |

### Global sampling of CDR1 sequence diversity using the LOGAN database

The LOGAN database^68^ was used to download from contigs from accessions listed as *Candida albicans* which mapped to our Cdr1 reference sequence (Uniprot accession: XP_723209) as per their tutorial on Github (https://github.com/IndexThePlanet/Logan/blob/main/Chickens.md), resulting in 5362 sequence. We found that a further 799 accessions had been deposited since the latest LOGAN timestamp, so these were downloaded from AWS and assembled manually using minimap2^70^. After assembling and aligning to the reference sequence, and gaps were filled with the corresponding nucleotides of the reference sequence to create complete sequences for downstream analysis. This resulted in 6161 sequences that were then analysed for duplicates, reducing the dataset to 511 unique sequences. These contained 106 evolutionarily informative sites that were then analysed using sequence logo^69^ (see Figure 6).

---

**Supplementary Figure 1.**
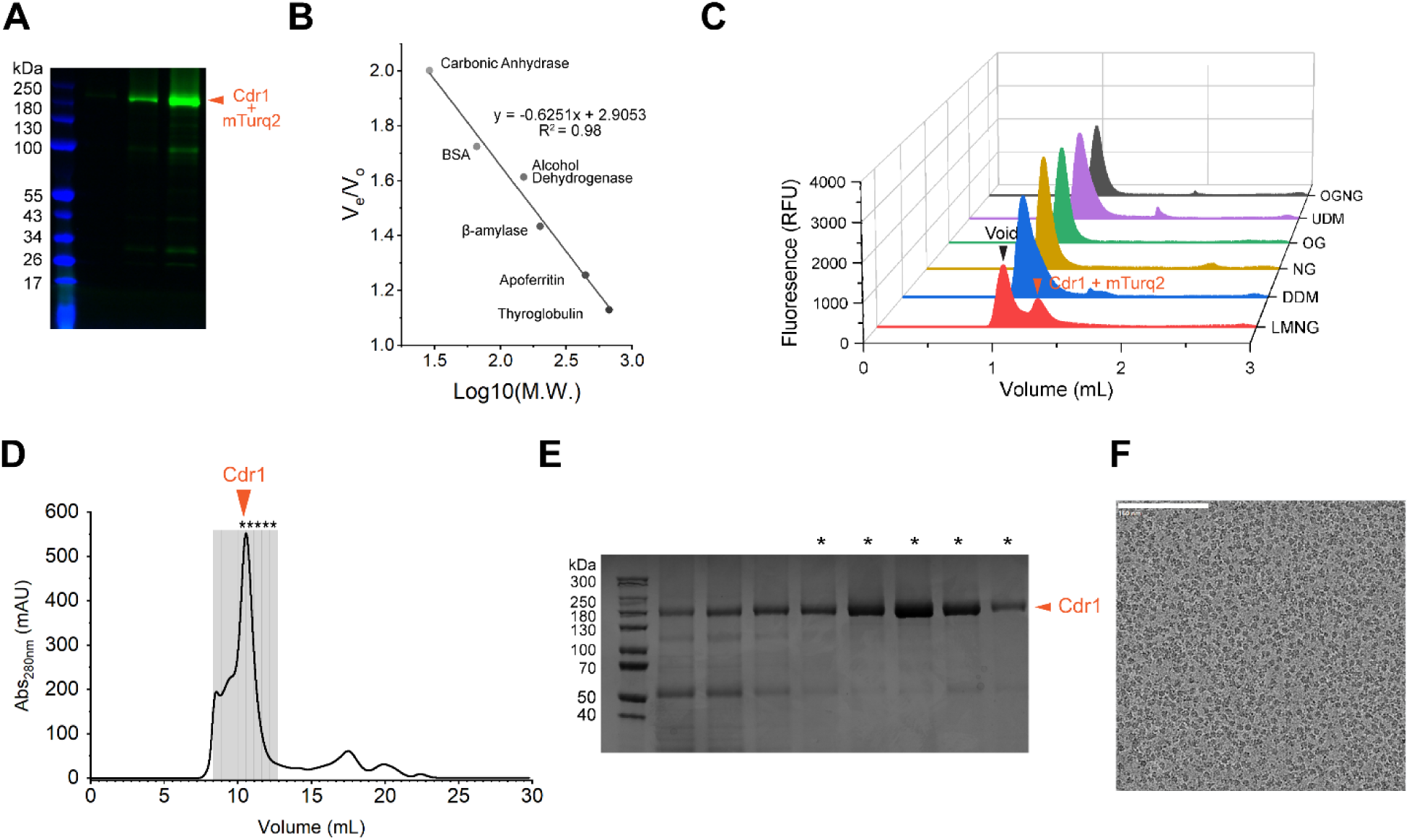
Detergent screening and purification of Cdr1. **(A)** In-gel fluorescence of first three elution fractions of strep-tatin purified mTurquoise2 fusion Cdr1. **(B)** Molecular weight calibration curve using the elution positions of the standards. V_e_ and V_o_ denotes protein retention volume and void volume, respectively. **(C)** FSEC profiles of detergent solubilised mTurquoise2 fusion Cdr1. **(D)** SEC profile of post Cdr1 reconstituted in LMNG. Grey bars indicate individual fractions taken for SDS-PAGE. **(E)** SDS-PAGE of SEC fractions. Asterisks indicate fractions used for downstream experiments. **(F)** Representative micrograph of Cdr1-LMNG.

**Supplementary Figure 2.**
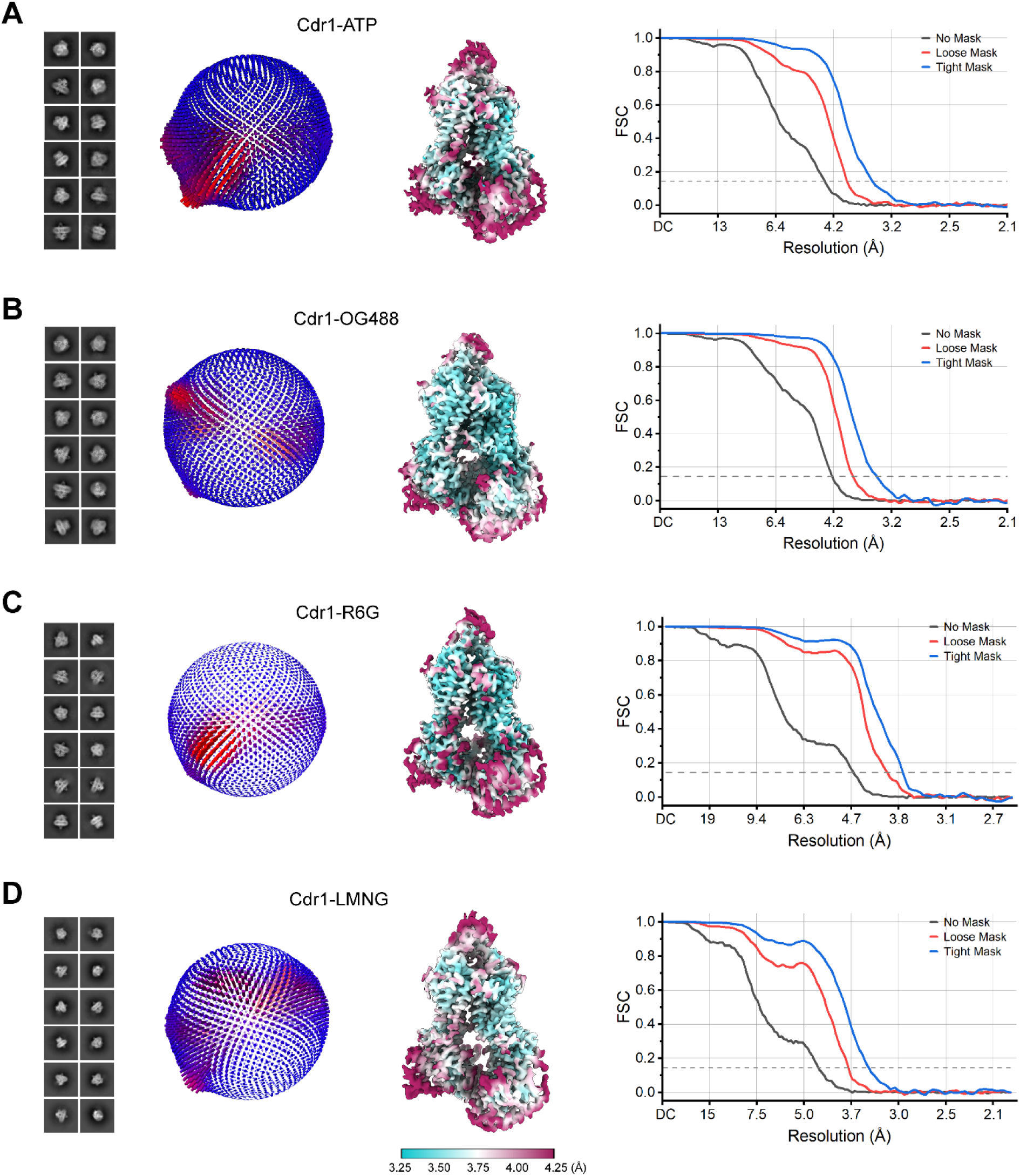
Cryo-EM data processing of Cdr1. **(A)** Cdr1-ATP (σ = 4.0) . **(B)** Cdr1-OG488 (σ = 2.5). **(C)** Cdr1-R6G (σ = 6.2). **(D)** Cdr1-LMNG (σ = 6.0). 2D classes, angular distribution, local resolution estimation of sharpened map prior to density modified with Phenix and Fourier shell correlation curve are shown in order from left to right.

**Supplementary Figure 3.**
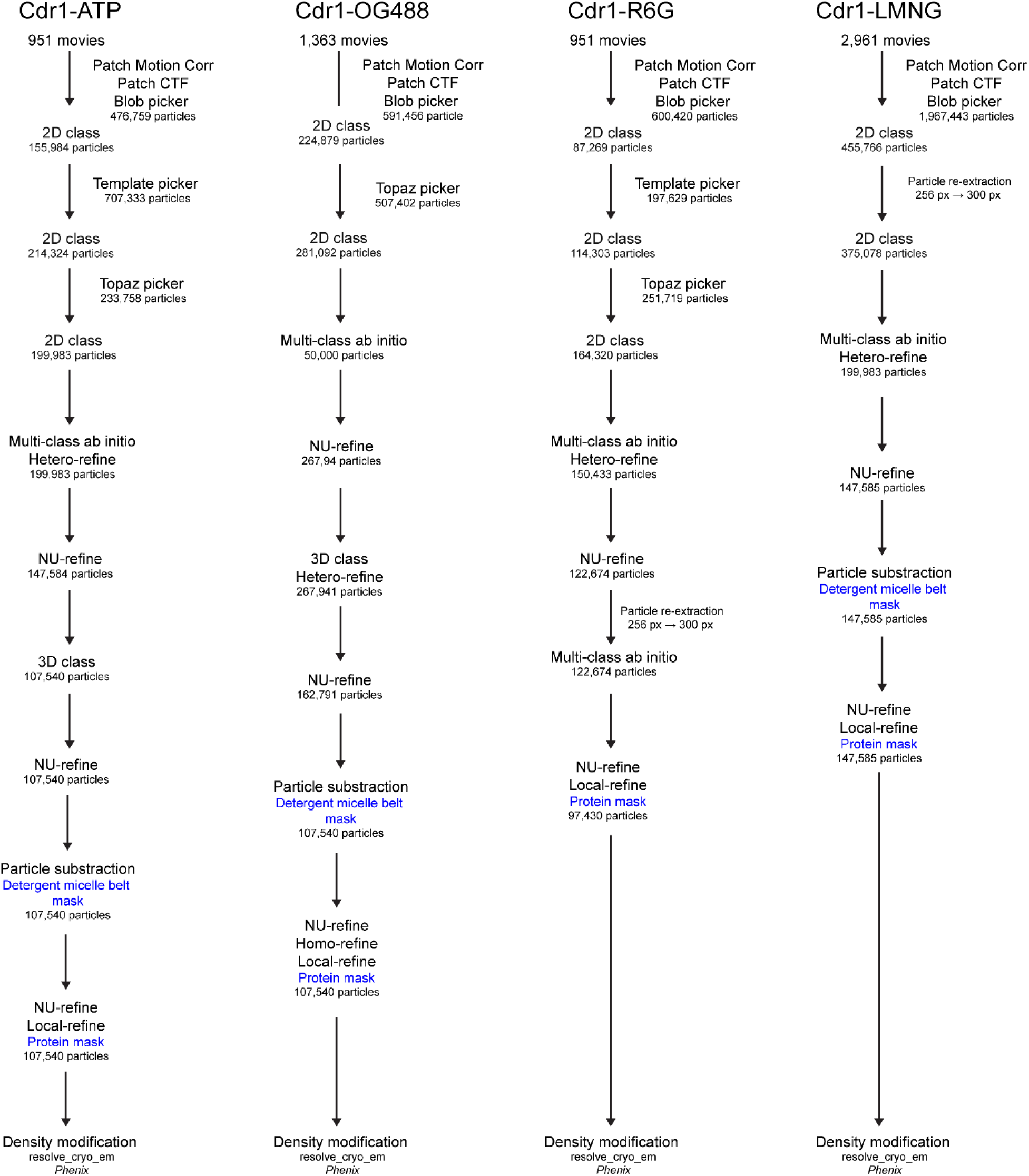
Cryo-EM data processing pipeline for each data set.

**Supplementary Figure 4.**
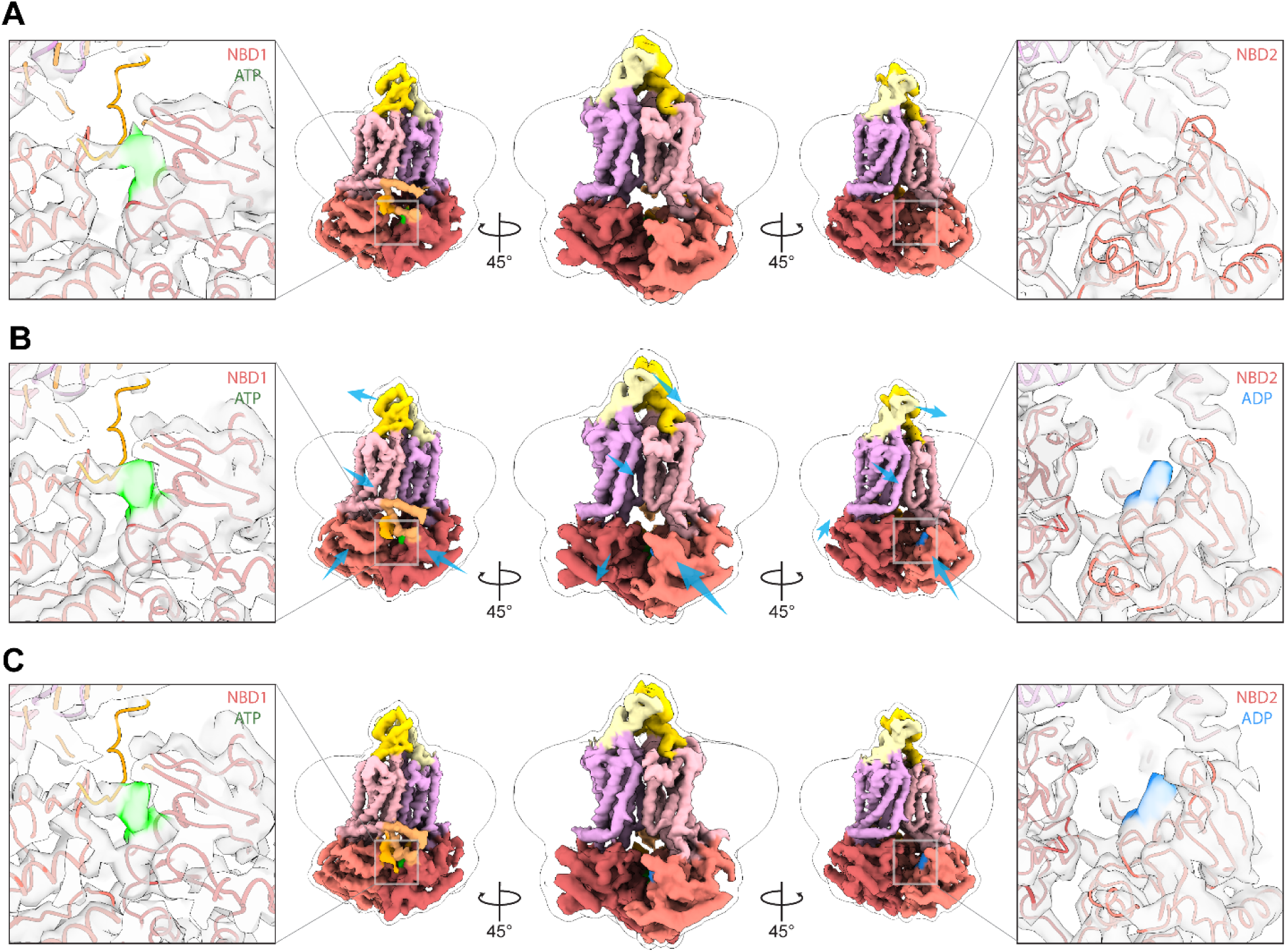
3D variability analysis of Cdr1-ATP using cryoSPARC v4.1.0^57^. Three Cryo-EM density maps representing distinct stages of the volume series are depicted: (A) the initial, (B) the intermediate, and (E) the final volume. Left and right panels of each figure display the surface representation of Cryo-EM density overlaid on the nucleotide-binding domains (NBDs) of the Cdr1-ATP model, shown in ribbon representation. The Cryo-EM densities for ATP and ADP are highlighted in lime green and blue, respectively. The density maps are color-coded to illustrate domain organization, while the detergent micelle belt is represented by a 3σ Gaussian-filtered map overlaying Cdr1-ATP. Blue arrows indicate the direction of map movement throughout the volume series. Map series contour at σ = 6.0 was used.

**Supplementary Figure 5.**
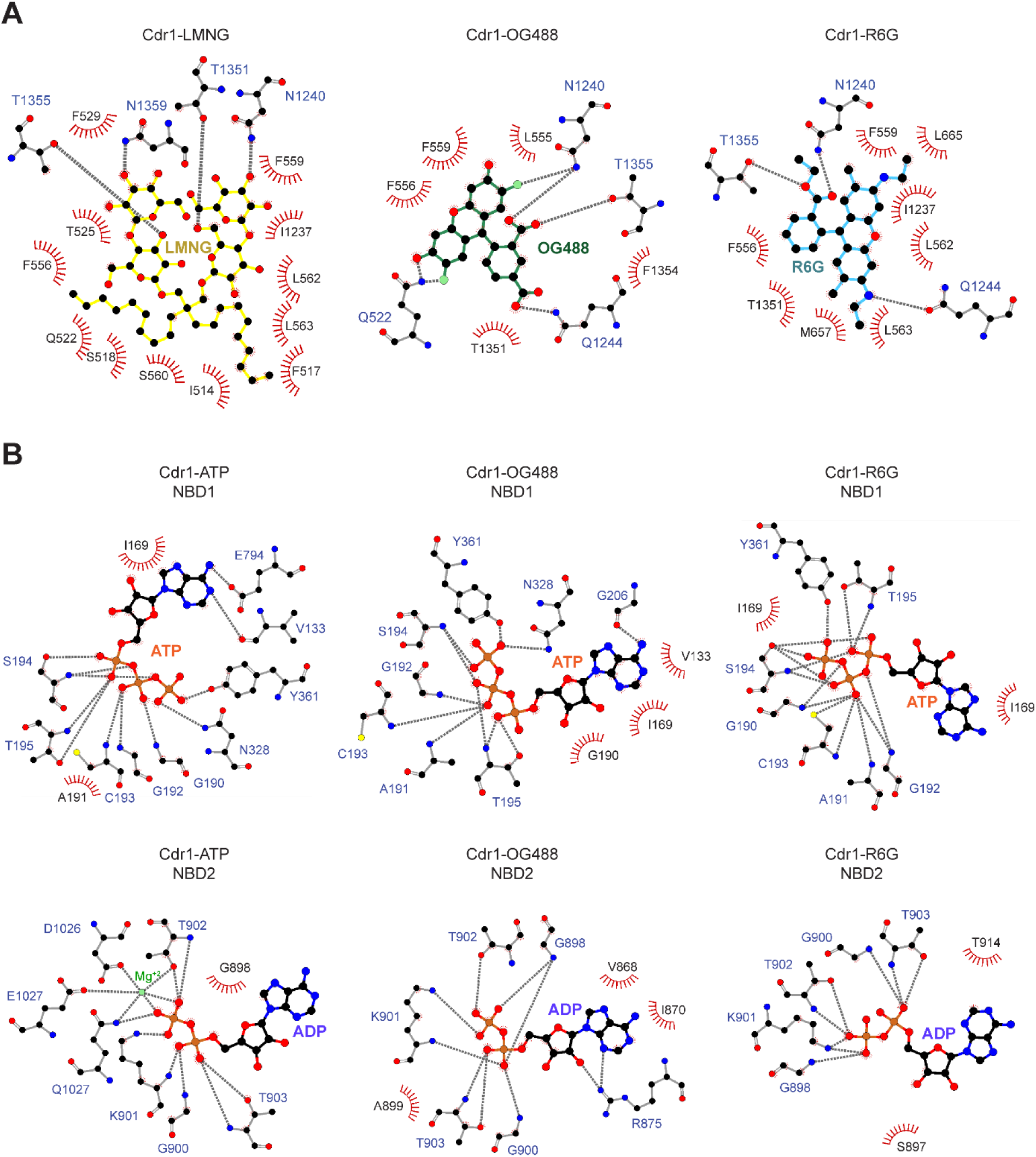
Protein-ligand interaction profile within Cdr1. LigPlot+2-based schematic showing molecular interactions between ligands, nucleotides and metal with Cdr1. **(A)** Ligand-protein interactions within the TMD binding site of Cdr1. **(B)** Nucleotide and metal ion interactions with NBDs of Cdr1. Residues participating in hydrogen bond, LMNG, OG488, R6G and nucleotides are depicted in stick representation. Bristled crescents indicate hydrophobic interactions and hydrogen bonds are shown as grey dashed lines. Atom colour key: C = black, N = blue, O = red, P = orange, F = lime green, Mg^2+^ = green.

**Supplementary Figure 6.**
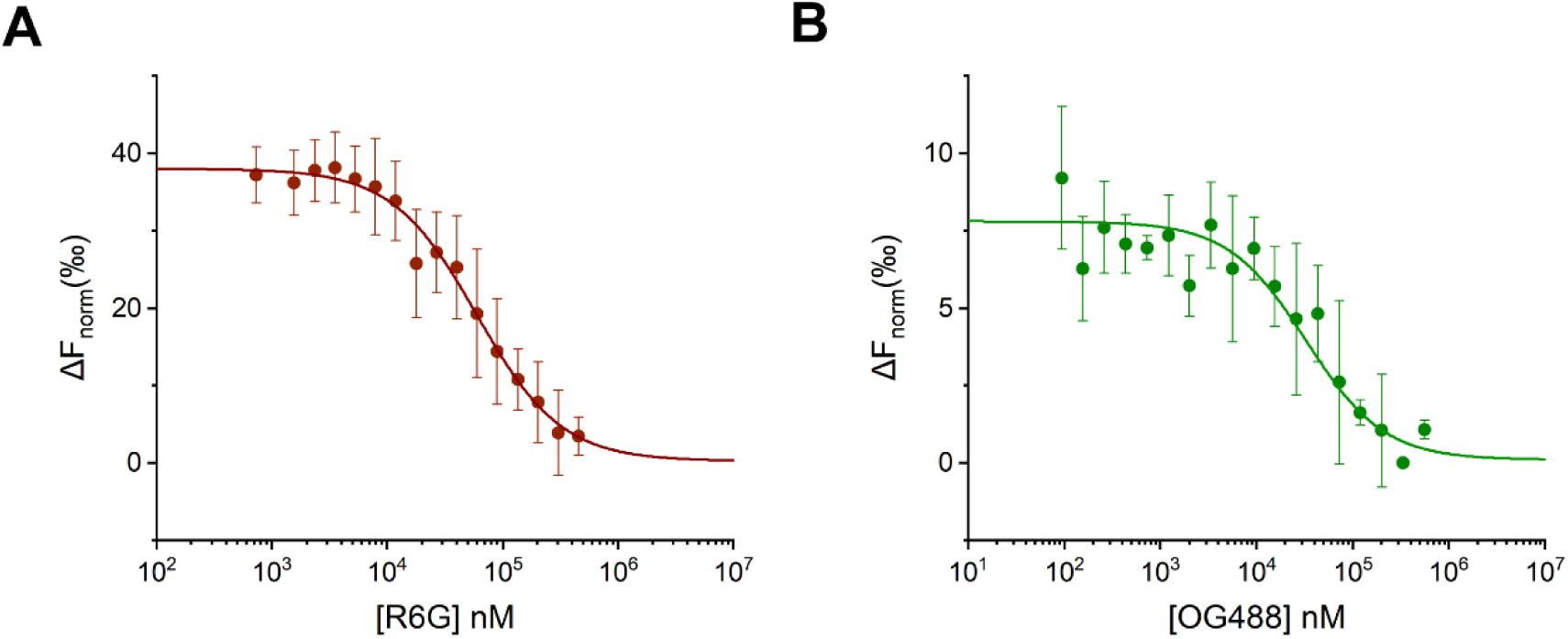
Microscale thermophoresis (MST) binding assay. **(A)** Binding interaction between R6G and Cdr1**. (B)** Interaction between OG488 and Cdr1. Error bars represent standard deviations from three experimental repeats.

**Supplementary Figure 7.**
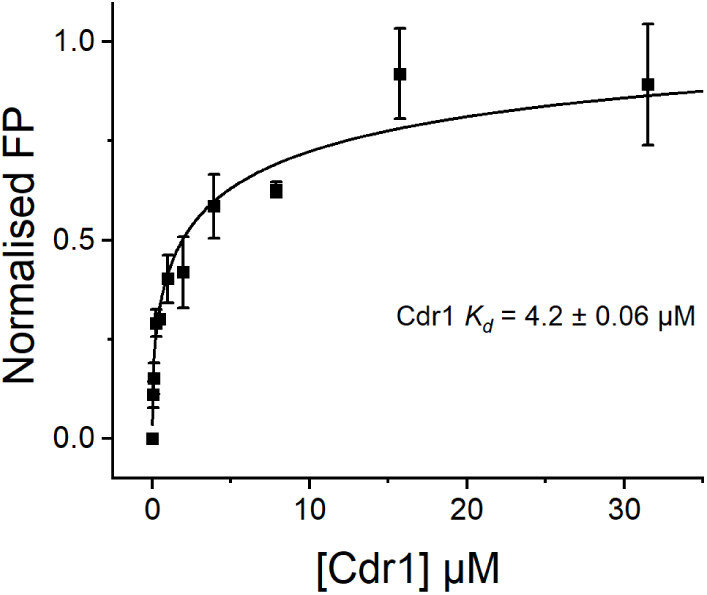
Fluorescence polarisation (FP) of direct binding of OG488 to Cdr1. The concentration of OG488 used was at 1 µM. Error bars represent standard deviations from two individual repeat measurements.

**Supplementary Figure 8.**
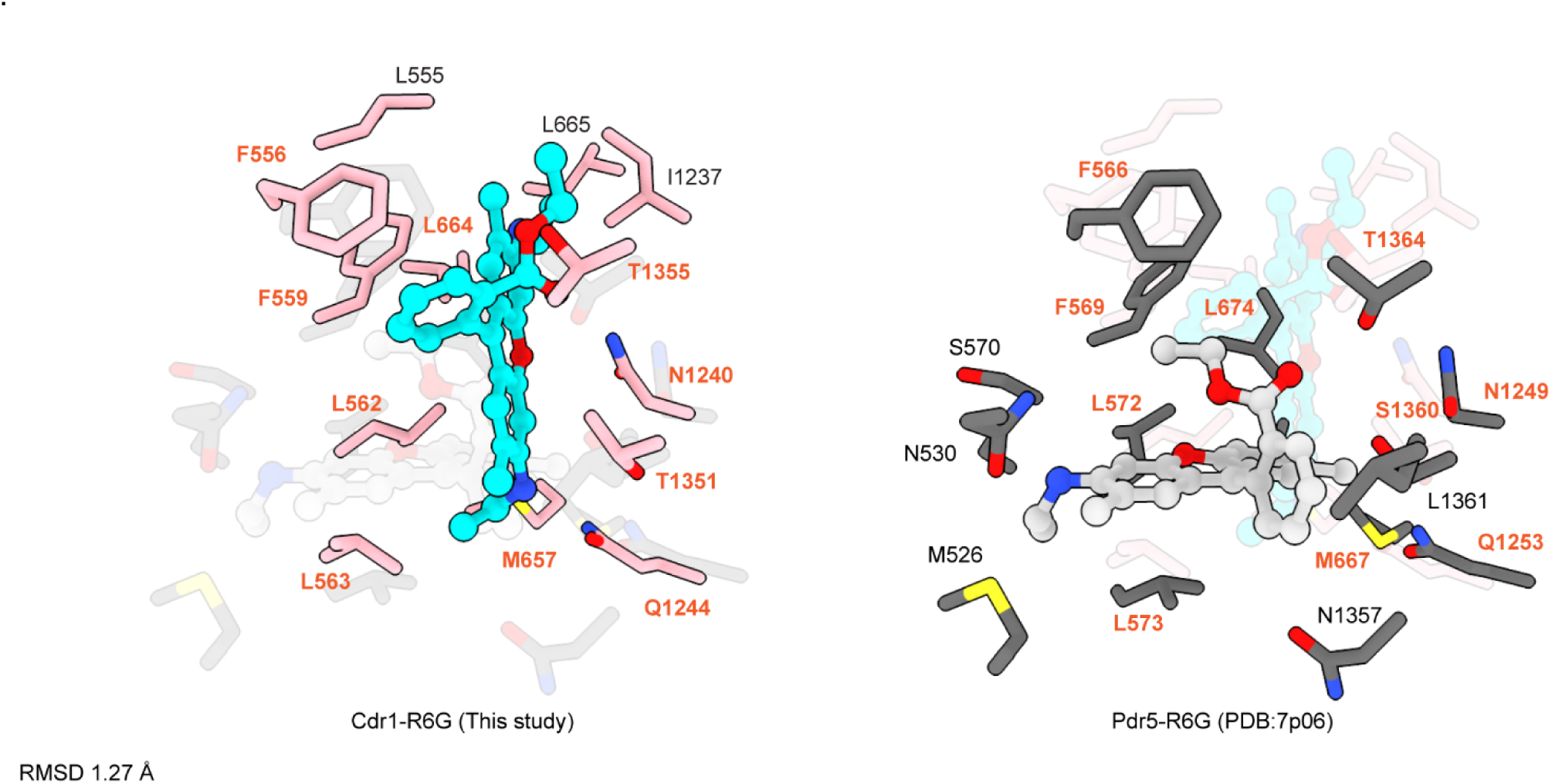
Structural alignment of Cdr1-R6G and R6G bound Pdr5 (PDB:7p06) reveals a 1.27 Å RMSD, highlighting ligand-binding residues within 4 Å distance range from the R6G and marking equivalents in orange.

**Supplementary Table1.** Cryo-EM data collection, refinement, and validation statistics.

|  | Cdr1-ATP | Cdr1-LMNG | Cdr1-OG488 | Cdr-R6G |
| --- | --- | --- | --- | --- |
| **Data collection and processing** |  |  |  |  |
| Magnification | 120,000 | 100,000 | 120,000 | 120,000 |
| Voltage (kV) | 200 | 200 | 200 | 200 |
| Electron exposure (e-/Å<sup>2</sup>) | 40 | 40 | 40 | 40 |
| Defocus range (μm) | -0.5 to -2.0 | -0.5 to -2.7 | -0.5 to -2.0 | -0.5 to -2.0 |
| Pixel size (Å) | 0.996 | 1.2512 | 0.996 | 0.996 |
| Symmetry imposed | C1 | C1 | C1 | C1 |
| Initial particle images (no.) | 476,759 | 431,140 | 620,086 | 600,420 |
| Final particle images (no.) | 107,540 | 147,585 | 162,791 | 97,430 |
| Map resolution (Å) | 3.50 | 3.73 | 3.49 | 3.45 |
| FSC threshold | 0.143 | 0.143 | 0.143 | 0.143 |
| **Refinement** |  |  |  |  |
| Initial model used (PDB code) | Cdr1-OG488 | AlphaFold model | AlphaFold model | Cdr1-OG488 |
| FSC (model) threshold | 0.5 | 0.5 | 0.5 | 0.5 |
| Model composition |  |  |  |  |
| Non-hydrogen atoms | 11,051 |  |  |  |
| Protein residues | 1373 |  |  |  |
| Ligands | MG: 1<br>ADP: 1<br>ATP: 1 | LMN: 1 | ORE: 1<br>ADP: 1<br>ATP: 1 | RQH: 1<br>ADP: 1<br>ATP: 1 |
| B factors (Å<sup>2</sup>) |  |  |  |  |
| Protein | 69.13/154.52/95 | 56.76/178.42/95.02 | 96.32/213.26/141.34 | 96.32/213.26/141.34 |
| Ligand | 96.69/121.72/1024 | 20.00/20.00/20.00 | 145.91/165.41/152.91 | 145.91/165.41/152.91 |
| R.M.S. deviations |  |  |  |  |
| Bond lengths (Å) | 0.014 | 0.014 | 0.012 | 0.013 |
| Bond angles (°) | 1.908 | 1.895 | 1.614 | 1.751 |
| **Validation** |  |  |  |  |
| MolProbity score | 0.54 | 0.50 | 0.70 | 0.52 |
| Clashscore | 0.09 | 0.00 | 0.61 | 0.05 |
| Poor rotamers (%) | 0.09 | 0.00 | 0.26 | 0.26 |
| **Ramachandran plot** |  |  |  |  |
| Favored (%) | 98.24 | 99.55 | 98.73 | 98.58 |
| Allowed (%) | 1.68 | 0.45 | 1.27 | 1.42 |
| Disallowed (%) | 0.07 | 0.00 | 0.00 | 0.00 |

**Supplementary Table 2.**
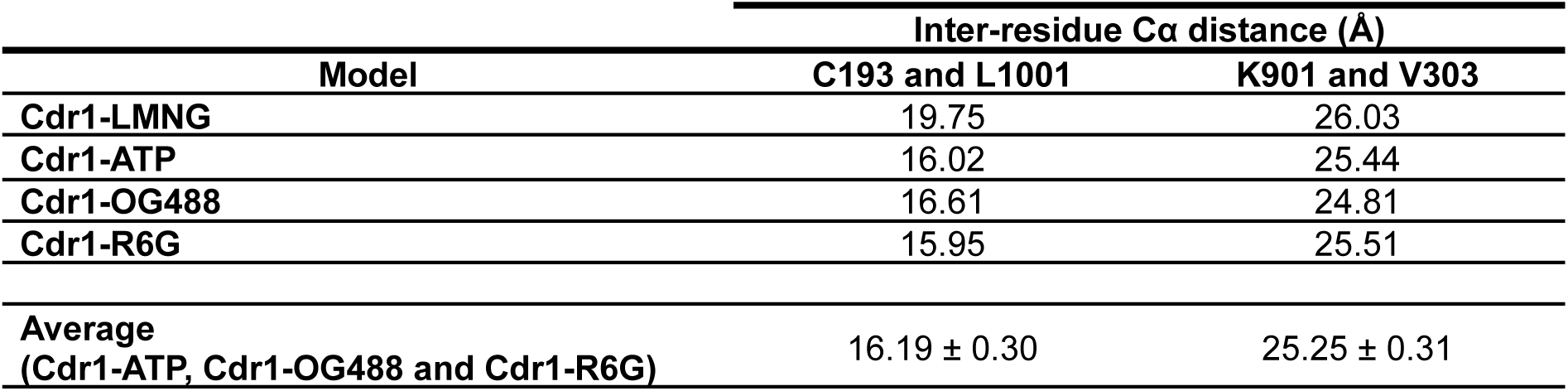
Inter-residue Cα distance measurements between NBDs.

| Model | Inter-residue Cα distance (Å) | |
| --- | --- | --- |
|  | C193 and L1001 | K901 and V303 |
| Cdr1-LMNG | 19.75 | 26.03 |
| Cdr1-ATP | 16.02 | 25.44 |
| Cdr1-OG488 | 16.61 | 24.81 |
| Cdr1-R6G | 15.95 | 25.51 |
| Average<br>(Cdr1-ATP, Cdr1-OG488 and Cdr1-R6G) | 16.19 ± 0.30 | 25.25 ± 0.31 |

**Supplementary Table 3.** Structural alignment of Cdr1.

| Reference model | Model aligned | RMSD<sub>across all pairs</sub> (Å) |
| --- | --- | --- |
| Cdr1-LMNG | Cdr1-ATP | 3.900 |
|  | Cdr1-OG488 | 3.042 |
|  | Cdr1-R6G | 2.416 |
| Cdr1-ATP | Cdr1-OG488 | 1.643 |
|  | Cdr1-R6G | 1.688 |

**Supplementary Table 4.** Kd values of R6G and OG488 binding to detergent reconstituted Cdr1 determined by MST.

|  |  |
| --- | --- |
| **R6G** | **$59.75 \pm 7.1 \mu\text{M}$** |
| **OG488** | **$32.798 \pm 8.2 \mu\text{M}$** |

## Notes

### Competing Interest Statement

The authors have declared no competing interest.

## References

1. Centers for Disease Control and Prevention (CDC). *Fungal Diseases and COVID-19. Centers for Disease Control and Prevention*. https://www.cdc.gov/fungal/hcp/covid-fungal/index.html (2021).

2. Forsberg, K. et al. Candida auris: The recent emergence of a multidrug-resistant fungal pathogen. Med. Mycol. 57, 1–12 (2019).

3. Castanheira, M., Deshpande, L. M., Davis, A. P., Rhomberg, P. R. & Pfaller, M. A. Monitoring Antifungal Resistance in a Global Collection of Invasive Yeasts and Molds: Application of CLSI Epidemiological Cutoff Values and Whole-Genome Sequencing Analysis for Detection of Azole Resistance in Candida albicans. Antimicrob. Agents Chemother. 61, e00906–17 (2017).

4. Esfahani, A. et al. Up-regulation of CDR1 and MDR1 efflux pump genes and fluconazole resistance are involved in recurrence in Candida albicans-induced vulvovaginal candidiasis. Diagn. Microbiol. Infect. Dis. 109, 116242 (2024).

5. Kean, R. & Ramage, G. Combined Antifungal Resistance and Biofilm Tolerance: the Global Threat of Candida auris. mSphere 4, (2019).

6. Kainz, K., Bauer, M. A., Madeo, F. & Carmona-Gutierrez, D. Fungal infections in humans: The silent crisis. Microb. Cell 7, 143–145 (2020).

7. World Health Organization. WHO fungal priority pathogens list to guide research, development and public health action. (2022).

8. Denning, D. W. Global incidence and mortality of severe fungal disease. Lancet Infect. Dis. 24, e428–e438 (2024).

9. Lee, Y., Robbins, N. & Cowen, L. E. Molecular mechanisms governing antifungal drug resistance. Npj Antimicrob. Resist. 1, 5 (2023).

10. Li, Q. et al. Sterol uptake and sterol biosynthesis act coordinately to mediate antifungal resistance in Candidaï¿½glabrata under azole and hypoxic stress. Mol. Med. Rep. (2018) doi:10.3892/mmr.2018.8716.

11. Rybak, J. M. et al. Mutations in TAC1B : a Novel Genetic Determinant of Clinical Fluconazole Resistance in Candida auris. mBio 11, 395–410 (2020).

12. Gao, J. et al. Candida albicans gains azole resistance by altering sphingolipid composition. Nat. Commun. 9, 4495 (2018).

13. Rybak, J. M. et al. Abrogation of Triazole Resistance upon Deletion of CDR1 in a Clinical Isolate of Candida auris. Antimicrob. Agents Chemother. 63, 1–7 (2019).

14. Thomas, C. et al. Structural and functional diversity calls for a new classification of ABC transporters. FEBS Lett. 594, 3767–3775 (2020).

15. Banerjee, A. et al. Structure, function, and inhibition of catalytically asymmetric ABC transporters: Lessons from the PDR subfamily. Drug Resist. Updat. 71, 100992 (2023).

16. Prasad, R., Banerjee, A., Khandelwa, N. K. & Dhamgaye, S. The ABCs of Candida albicans multidrug transporter Cdr1. Eukaryot. Cell 14, 1154–1164 (2015).

17. Smriti et al. ABC transporters CdrLp, Cdr2p and Cdr3p of a human pathogen candida albicans are general phospholipid translocators. Yeast 19, 303–318 (2002).

18. Baghel, P. et al. Multidrug ABC transporter Cdr1 of Candida albicans harbors specific and overlapping binding sites for human steroid hormones transport. Biochim. Biophys. Acta - Biomembr. 1859, 1778–1789 (2017).

19. Golin, J. & Schmitt, L. Pdr5: A master of asymmetry. Drug Resist. Updat. 71, 101010 (2023).

20. Godinho, C. P. et al. Pdr18 is involved in yeast response to acetic acid stress counteracting the decrease of plasma membrane ergosterol content and order. Sci. Rep. 8, 7860 (2018).

21. Sun, Y. & Li, X. Cholesterol efflux mechanism revealed by structural analysis of human ABCA1 conformational states. *Nat*. Cardiovasc. Res. 1, 238–245 (2022).

22. Le, L. T. M. et al. Cryo-EM structures of human ABCA7 provide insights into its phospholipid translocation mechanisms. EMBO J. 42, e111065 (2023).

23. Xie, T., Zhang, Z., Fang, Q., Du, B. & Gong, X. Structural basis of substrate recognition and translocation by human ABCA4. Nat. Commun. 12, 3853 (2021).

24. Shukla, S., et al. *Candida* Drug Resistance Protein 1, a Major Multidrug ATP Binding Cassette Transporter of *Candida albicans*, Translocates Fluorescent Phospholipids in a Reconstituted System. Biochemistry 46, 12081–12090 (2007).

25. Lamping, E. et al. Fungal PDR transporters: Phylogeny, topology, motifs and function. Fungal Genet. Biol. 47, 127–142 (2010).

26. Harris, A. et al. Structure and efflux mechanism of the yeast pleiotropic drug resistance transporter Pdr5. Nat. Commun. 12, 5254 (2021).

27. Raschka, S. L., Harris, A., Luisi, B. F. & Schmitt, L. Flipping and other astonishing transporter dance moves in fungal drug resistance. BioEssays 44, 2200035 (2022).

28. Kovalchuk, A. & Driessen, A. J. Phylogenetic analysis of fungal ABC transporters. BMC Genomics 11, 177 (2010).

29. Peng, Y. et al. Cryo-EM structures of Candida albicans Cdr1 reveal azole-substrate recognition and inhibitor blocking mechanisms. Nat. Commun. 15, 7722 (2024).

30. Niimi, M., Niimi, K., Tanabe, K., Cannon, R. D. & Lamping, E. Inhibitor-Resistant Mutants Give Important Insights into Candida albicans ABC Transporter Cdr1 Substrate Specificity and Help Elucidate Efflux Pump Inhibition. Antimicrob. Agents Chemother. 66, (2022).

31. Tanabe, K. et al. FK506 Resistance of *Saccharomyces cerevisiae* Pdr5 and *Candida albicans* Cdr1 Involves Mutations in the Transmembrane Domains and Extracellular Loops. Antimicrob. Agents Chemother. 63, e01146–18 (2019).

32. Coste, A. et al. A Mutation in Tac1p, a Transcription Factor Regulating *CDR1* and *CDR2*, Is Coupled With Loss of Heterozygosity at Chromosome 5 to Mediate Antifungal Resistance in *Candida albicans*. Genetics 172, 2139–2156 (2006).

33. Ropars, J. et al. Gene flow contributes to diversification of the major fungal pathogen Candida albicans. Nat. Commun. 9, 2253 (2018).

34. Mario-Vasquez, J. E. et al. Finding Candida auris in public metagenomic repositories. PLOS ONE 19, e0291406 (2024).

35. Chikhi, R., Raffestin, B., Korobeynikov, A., Edgar, R. & Babaian, A. Logan: Planetary-Scale Genome Assembly Surveys Life’s Diversity. Preprint at 10.1101/2024.07.30.605881 (2024).

36. Nakamura, K. et al. Functional Expression of *Candida albicans* Drug Efflux Pump Cdr1p in a *Saccharomyces cerevisiae* Strain Deficient in Membrane Transporters. Antimicrob. Agents Chemother. 45, 3366–3374 (2001).

37. Banerjee, A. et al. Cdr1p highlights the role of the non-hydrolytic ATP-binding site in driving drug translocation in asymmetric ABC pumps. Biochim. Biophys. Acta BBA - Biomembr. 1862, 183131 (2020).

38. Li, S. et al. The δ subunit of F1Fo-ATP synthase is required for pathogenicity of Candida albicans. Nat. Commun. 12, 6041 (2021).

39. Pan, X., Wang, H., Li, C., Zhang, J. Z. H. & Ji, C. MolGpka: A Web Server for Small Molecule p *K* _a_ Prediction Using a Graph-Convolutional Neural Network. J. Chem. Inf. Model. 61, 3159–3165 (2021).

40. Ashkenazy, H. et al. ConSurf 2016: an improved methodology to estimate and visualize evolutionary conservation in macromolecules. Nucleic Acids Res. 44, W344–W350 (2016).

41. Shaner, M. C., Blair, I. M. & Schneider, T. D. Sequence logos: a powerful, yet simple, tool. in [1993] Proceedings of the Twenty-sixth Hawaii International Conference on System Sciences vol. i 813–821 vol.1 (1993).

42. Ernst, R. et al. A mutation of the H-loop selectively affects rhodamine transport by the yeast multidrug ABC transporter Pdr5. Proc. Natl. Acad. Sci. 105, 5069–5074 (2008).

43. Drew, D., Newstead, S., Sonoda, Y., Kim, H. & Heijne, G. V. GFP-based optimization scheme for the overexpression and purification of eukaryotic membrane proteins in Saccharomyces cerevisiae. Nat. Protoc. 3, 784–798 (2009).

44. Newstead, S., Kim, H., von Heijne, G., Iwata, S. & Drew, D. High-throughput fluorescent-based optimization of eukaryotic membrane protein overexpression and purification in Saccharomyces cerevisiae. Proc. Natl. Acad. Sci. 104, 13936–13941 (2007).

45. Lee, M. E., DeLoache, W. C., Cervantes, B. & Dueber, J. E. A Highly Characterized Yeast Toolkit for Modular, Multipart Assembly. ACS Synth. Biol. 4, 975–986 (2015).

46. Mastronarde, D. N. Automated electron microscope tomography using robust prediction of specimen movements. J. Struct. Biol. 152, 36–51 (2005).

47. Punjani, A., Rubinstein, J. L., Fleet, D. J. & Brubaker, M. A. CryoSPARC: Algorithms for rapid unsupervised cryo-EM structure determination. Nat. Methods 14, 290–296 (2017).

48. Bepler, T. et al. Positive-unlabeled convolutional neural networks for particle picking in cryo-electron micrographs. Nat. Methods 16, 1153–1160 (2019).

49. Pettersen, E. F. et al. UCSF ChimeraX : Structure visualization for researchers, educators, and developers. Protein Sci. 30, 70–82 (2021).

50. Terwilliger, T. C., Ludtke, S. J., Read, R. J., Adams, P. D. & Afonine, P. V. Improvement of cryo-EM maps by density modification. Nat. Methods 17, 923–927 (2020).

51. Adams, P. D. et al. PHENIX: A comprehensive Python-based system for macromolecular structure solution. Acta Crystallogr. D Biol. Crystallogr. 66, 213–221 (2010).

52. Jumper, J. et al. Highly accurate protein structure prediction with AlphaFold. Nature 596, 583–589 (2021).

53. Afonine, P. V. et al. Real-space refinement in *PHENIX* for cryo-EM and crystallography. Acta Crystallogr. Sect. Struct. Biol. 74, 531–544 (2018).

54. Emsley, P. & Cowtan, K. Coot: Model-building tools for molecular graphics. Acta Crystallogr. D Biol. Crystallogr. 60, 2126–2132 (2004).

55. Croll, T. I. ISOLDE: a physically realistic environment for model building into low-resolution electron-density maps. Acta Crystallogr. Sect. Struct. Biol. 74, 519–530 (2018).

56. Moriarty, N. W., Grosse-Kunstleve, R. W. & Adams, P. D. *electronic Ligand Builder and Optimization Workbench* (*eLBOW*): a tool for ligand coordinate and restraint generation. Acta Crystallogr. D Biol. Crystallogr. 65, 1074–1080 (2009).

57. Davis, I. et al. MolProbity: all-atom contacts and structure validation for proteins and nucleic acids. Nucleic Acids Res 35, W375–83 (2007).

58. Trott, O. & Olson, A. J. AutoDock Vina: Improving the speed and accuracy of docking with a new scoring function, efficient optimization, and multithreading. J. Comput. Chem. 31, 455–461 (2010).

59. Jo, S., Lim, J. B., Klauda, J. B. & Im, W. CHARMM-GUI Membrane Builder for Mixed Bilayers and Its Application to Yeast Membranes. Biophys. J. 96, 50–58 (2009).

60. Lomize, M. A., Pogozheva, I. D., Joo, H., Mosberg, H. I. & Lomize, A. L. OPM database and PPM web server: resources for positioning of proteins in membranes. Nucleic Acids Res. 40, D370–D376 (2012).

61. Van Der Spoel, D. et al. GROMACS: Fast, flexible, and free. J. Comput. Chem. 26, 1701–1718 (2005).

62. Vanommeslaeghe, K. & MacKerell, A. D. Automation of the CHARMM General Force Field (CGenFF) I: Bond Perception and Atom Typing. J. Chem. Inf. Model. 52, 3144–3154 (2012).

63. Huang, J. et al. CHARMM36m: an improved force field for folded and intrinsically disordered proteins. Nat. Methods 14, 71–73 (2017).

64. Hess, B., Bekker, H., Berendsen, H. J. C. & Fraaije, J. G. E. M. LINCS: A linear constraint solver for molecular simulations. J. Comput. Chem. 18, 1463–1472 (1997).

65. Darden, T., York, D. & Pedersen, L. Particle mesh Ewald: An *N* ⋅log(*N*) method for Ewald sums in large systems. J. Chem. Phys. 98, 10089–10092 (1993).

66. Verlet, L. Computer ‘Experiments’ on Classical Fluids. I. Thermodynamical Properties of Lennard-Jones Molecules. Phys. Rev. 159, 98–103 (1967).

67. Berendsen, H. J. C., Postma, J. P. M., Van Gunsteren, W. F., DiNola, A. & Haak, J. R. Molecular dynamics with coupling to an external bath. J. Chem. Phys. 81, 3684–3690 (1984).

68. Hoover, W. G. & Holian, B. L. Kinetic moments method for the canonical ensemble distribution. Phys. Lett. A 211, 253–257 (1996).

69. Parrinello, M. & Rahman, A. Polymorphic transitions in single crystals: A new molecular dynamics method. J. Appl. Phys. 52, 7182–7190 (1981).

70. Li, H. Minimap2: pairwise alignment for nucleotide sequences. Bioinformatics 34, 3094–3100 (2018).

